# Modeling uniquely human gene regulatory function in humanized mice

**DOI:** 10.1101/2019.12.11.873075

**Authors:** Emily V. Dutrow, Deena Emera, Kristina Yim, Severin Uebbing, Acadia A. Kocher, Martina Krenzer, Timothy Nottoli, Daniel B. Burkhardt, Smita Krishnaswamy, Angeliki Louvi, James P. Noonan

## Abstract

The evolution of uniquely human traits likely entailed changes in developmental gene regulation. Human Accelerated Regions (HARs), which include transcriptional enhancers harboring a significant excess of human-specific sequence changes, are leading candidates for driving gene regulatory modifications in human development. However, insight into whether HARs alter the level, distribution and timing of endogenous gene expression remains limited. We examined the role of the HAR *HACNS1* (HAR2) in human evolution by interrogating its molecular functions in a humanized mouse model. We find that *HACNS1* maintains its human-specific enhancer activity in humanized mice and that it modifies expression of *Gbx2*, which encodes a homeobox transcription factor, during limb development. Using single-cell RNA-sequencing, we demonstrate that *Gbx2* is upregulated in the chondrogenic mesenchyme of humanized limbs, supporting that *HACNS1* alters gene expression in cell types involved in skeletal patterning. Our findings illustrate that humanized mouse models provide mechanistic insight into how HARs modified gene expression in human evolution.

## Introduction

The evolution of uniquely human physical traits required human-specific genetic changes that altered development (*1, 2*). Discovering the locations of these changes in the genome and determining their biological impact is a major challenge. However, over the last decade comparative studies have begun to reveal potential genetic drivers underlying novel human biological features. These efforts have identified a prominent class of elements in the genome that are highly conserved across many species but show a significant excess of human-specific sequence changes (*3–7*). These elements, collectively named Human Accelerated Regions (HARs), are prime candidates to encode novel human molecular functions. Many HARs act as transcriptional enhancers during embryonic development, particularly in structures showing human-specific morphological changes such as the brain and limb (*7–12*). HARs have also been shown to exhibit human-specific changes in enhancer activity, both in transgenic assays and in massively parallel reporter assays in cultured cells (*7, 9, 10, 13–17*). These findings suggest a critical contribution for HARs in human evolution and support the long-standing hypothesis that changes in developmental gene regulatory programs contribute to evolutionary innovation (*18, 19*).

Despite these advances, the role of HARs in altering regulatory function *in vivo* remains poorly understood. We used a humanized mouse model approach to directly characterize the effects of human-specific sequence changes in HARs on gene expression and regulation during embryonic development (Fig. 1A). A similar genetic approach has been used to model the transcriptional and developmental effects of changes in enhancer activity in other mammalian lineages, notably bats (*20*). We chose to model *HACNS1* (also known as HAR2 or 2xHAR.3), as it exhibits the strongest acceleration signature of any noncoding HAR yet identified, with 13 human-specific substitutions in a 546 base-pair interval (Fig. 1B). *HACNS1* was also the first HAR demonstrated to exhibit a human-specific gain in enhancer activity during development. In a mouse transgenic enhancer assay, *HACNS1* was shown to drive increased expression of a *LacZ* reporter gene in the embryonic mouse limb compared to its chimpanzee and rhesus macaque orthologs (*10*). Furthermore, *HACNS1* exhibits increased levels of histone H3K27 acetylation (H3K27ac), which is correlated with enhancer activity, in human versus rhesus macaque and mouse embryonic limb (*11*). Together, these findings suggest that *HACNS1* may have contributed to changes in limb development during human evolution.

**Fig. 1.**
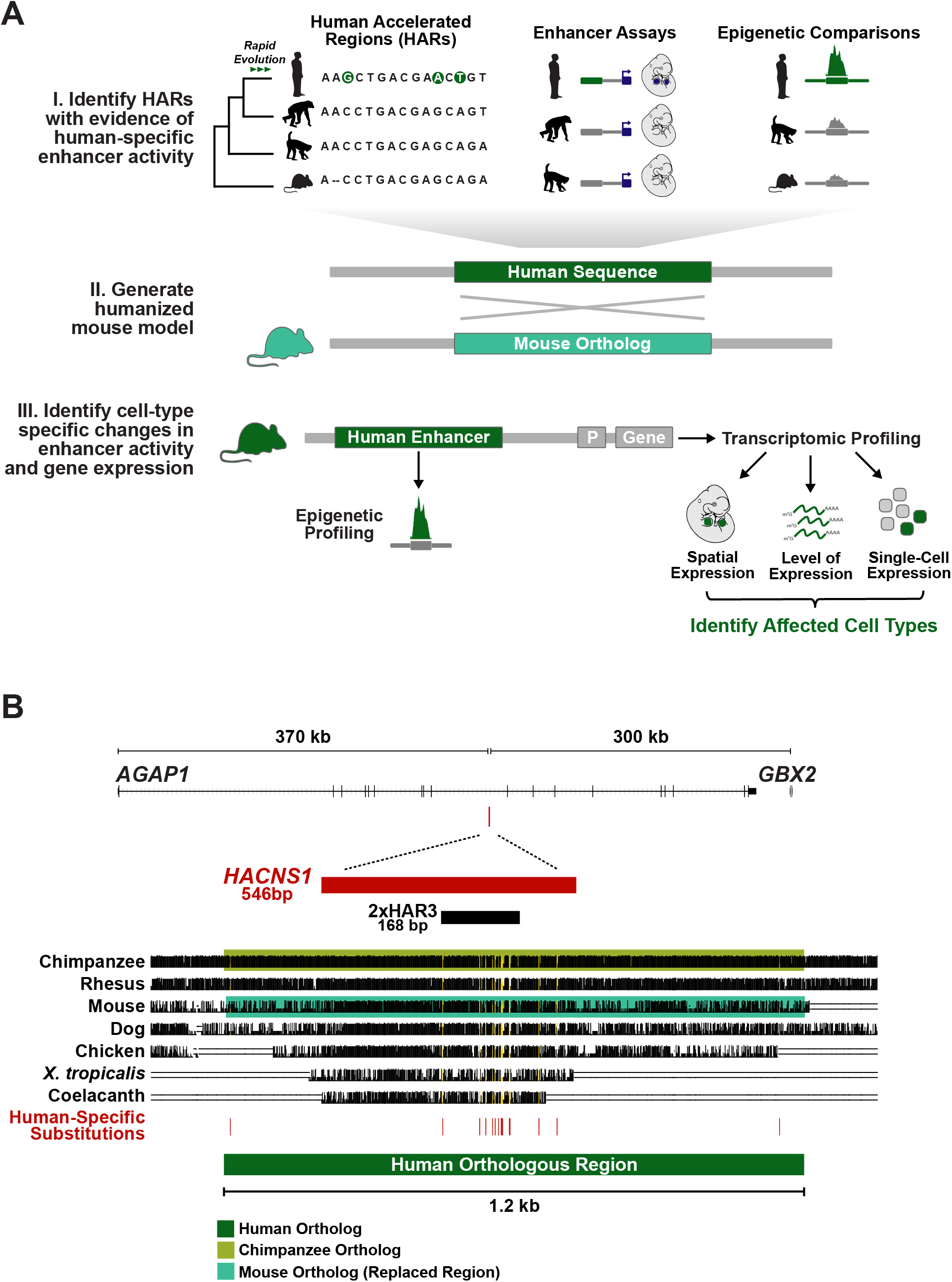
Generating a humanized mouse model for the Human Accelerated Region *HACNS1*. (**A**) Schematic illustrating the generalized workflow we developed to characterize the gene regulatory functions of HARs with prior evidence of human-specific enhancer activity, which we applied to *HACNS1* in this study. (**B**) The location of *HACNS1* in the human genome (GRCh37/hg19) relative to the nearby genes *AGAP1* and *GBX2*. Below, alignment of the human sequence used to generate the *HACNS1* humanized mouse with orthologous sequences from other vertebrate genomes, obtained from the UCSC hg19 100-way Multiz alignment (see Table S1 for coordinates). The chimpanzee orthologous sequence used to generate the chimpanzee control line is highlighted in olive, and the mouse sequence replaced in each line is highlighted in teal. The location of each human-specific substitution is indicated by a red line, and the corresponding positions in the alignment are highlighted in yellow. The locations of *HACNS1* and 2xHAR3 are shown above the alignment (*3, 6*).

To compare the functions of *HACNS1* and its chimpanzee and mouse orthologs in the same developmental system, we used homologous recombination to replace the endogenous mouse sequence with the human or chimpanzee counterpart. We found that *HACNS1* maintains its human-specific enhancer activity in the mouse embryo and alters the expression of the nearby transcription factor encoding gene *Gbx2* in limb chondrocytes, a cell type required for skeletal morphogenesis. Our findings support that HARs are capable of directing changes in endogenous gene expression during development and illustrate the power of humanized mouse models to provide insight into regulatory pathways and developmental mechanisms modified in human evolution.

## Results

### Generating an *HACNS1* humanized mouse model

We designed a targeting construct for homologous recombination including a 1.2 kb human sequence encompassing *HACNS1* that was previously shown to encode human-specific enhancer activity in transgenic mouse embryos (*10*). We replaced the orthologous mouse locus using homology-directed repair in C57BL6/J-*A^w-J^*/J (B6 agouti) embryonic stem (ES) cells (Fig. 1B, Fig. S1A,B, Table S1; Materials and Methods). To provide a control that would enable us to distinguish *bona fide* human-specific functions of *HACNS1* from possible primate-rodent differences, we used the same approach to generate a mouse model for the orthologous chimpanzee sequence (Fig S1A-C). The 1.2 kb chimpanzee sequence shows no evidence of evolutionary acceleration and includes 22 single nucleotide differences relative to the human sequence (*3*); fifteen of these differences are human-specific based on comparisons to other primate genomes (see Materials and Methods, Table S2). Previous studies indicate that multiple human-specific substitutions contribute to the gain of function in *HACNS1* (*10*). We found that 12 of the 15 substitutions introduced one or more predicted transcription factor binding sites that are specific to the human sequence (Table S2, Fig. S1A). An extensive comparison of sequence similarity and divergence among the human, chimpanzee, and mouse sequences is provided in the Supplemental Note (Supplementary Materials).

In order to verify the integrity of the edited loci, we sequenced a 40 kb region encompassing the human or chimpanzee sequence replacement, the homology arms used for targeting, and flanking genomic regions in mice homozygous for either *HACNS1*, or the chimpanzee ortholog (Fig. S1C; Materials and Methods). We found no evidence of aberrant editing, sequence rearrangements, or other off-target mutations at either edited locus. We also verified that each homozygous line carried two copies of the human or chimpanzee sequence using quantitative real-time PCR (qRT-PCR) (Fig. S1D).

### *HACNS1* exhibits human-specific enhancer activity in the humanized mouse embryo

We used chromatin immunoprecipitation (ChIP) to determine if *HACNS1* exhibits epigenetic signatures of increased enhancer activity in humanized mice. We first performed epigenetic profiling in the developing mouse limb bud based on prior evidence that *HACNS1* drives increased reporter gene activity in transgenic enhancer assays and exhibits increased H3K27ac marking in the human embryonic limb (*10, 11*). We profiled both H3K27ac and H3K4 dimethylation (H3K4me2), which is also associated with enhancer activity, in embryonic day (E) 11.5 limb buds from embryos homozygous for *HACNS1*, embryos homozygous for the chimpanzee ortholog, and wild type embryos. We found a strong signature of H3K27ac marking at *HACNS1* in the limb buds of *HACNS1* homozygous embryos (Fig. 2, S2). The chimpanzee and mouse sequences both showed significant but weaker H3K27ac enrichment relative to the human sequence, supporting the conclusion that *HACNS1* maintains its human-specific enhancer activity in the humanized mouse model.

**Fig. 2.**
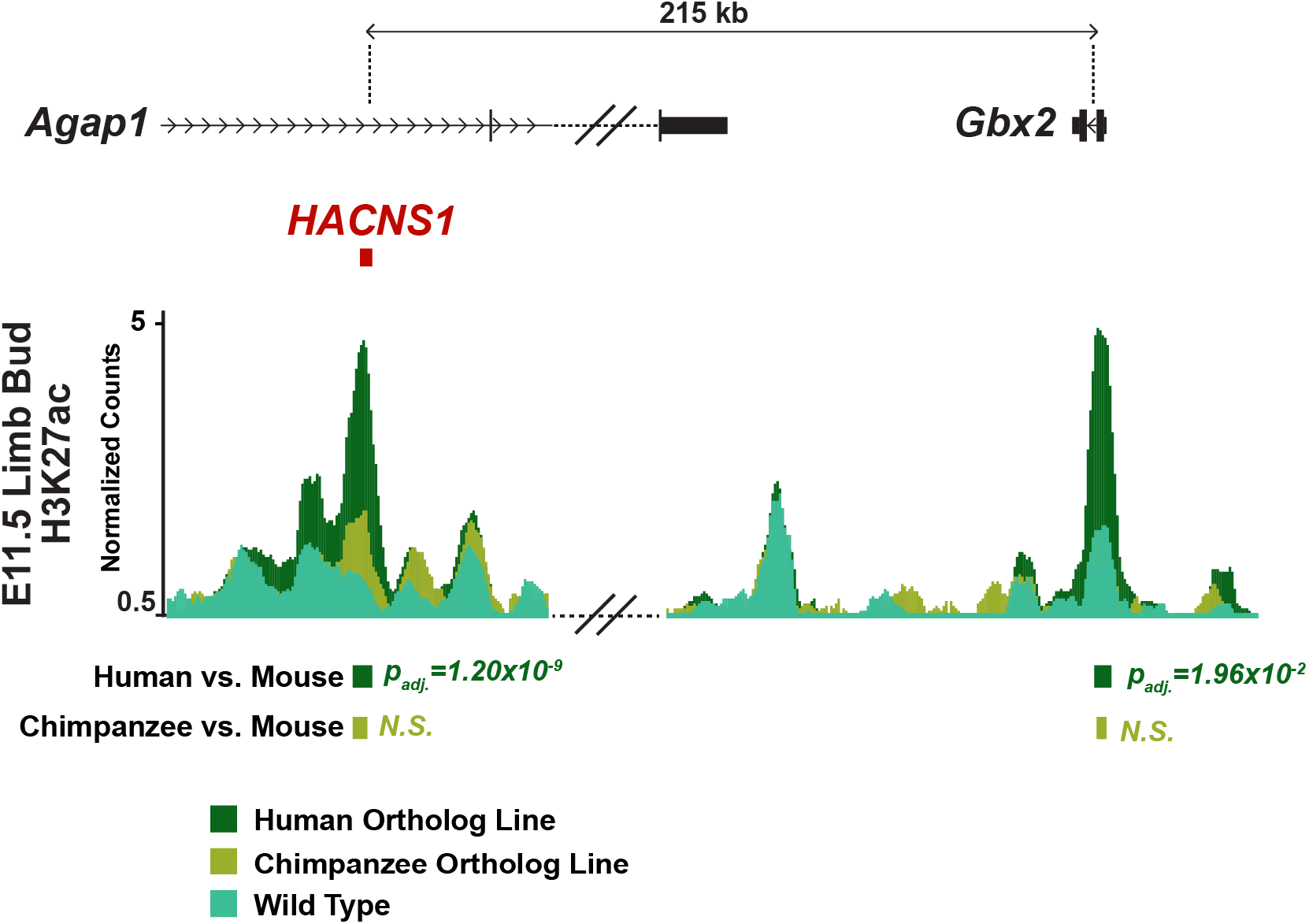
Epigenetic signatures of increased activity at *HACNS1* and the *Gbx2* promoter in the humanized mouse limb bud. Epigenetic profiling in humanized E11.5 limb bud compared to the chimpanzee ortholog line and wild type. The normalized H3K27ac signals are shown for the *HACNS1* line (in dark green), the chimpanzee ortholog line (in olive), and wild type (in teal) (see Materials and Methods). The location of the edited *HACNS1* locus in the humanized line relative to nearby genes is shown above the track. The double slanted lines indicate intervening H3K27ac signal data between the edited and wild type loci and *Gbx2* that were removed for clarity; see Fig. S2 for complete views for each line as well as input signals. H3K27ac peak calls showing significant increases in signal between *HACNS1* homozygous and wild type, and the corresponding peak regions compared between the chimpanzee control line and wild type, are shown below the signal track. Litter-matched embryos were used for each comparison (see Materials and Methods). *N.S.* = not significant. All peak calls for each line are shown in Fig. S2. Benjamini-Hochberg adjusted *P* values were obtained using DESeq2 implemented in HOMER (*21, 22*).

We used DESeq2 implemented in HOMER (see Materials and Methods) to identify genome-wide significant differences in H3K27ac and H3K4me2 levels in limb buds from mice homozygous for *HACNS1* or the chimpanzee ortholog versus wild type (*21, 22*). At the edited *HACNS1* locus, we found that H3K27ac and H3K4me2 levels were significantly increased in humanized limb compared to those at the endogenous mouse locus (Fig. 2, S2, Table S3). In contrast, the level of H3K27ac at the edited chimpanzee locus was not significantly different than that at the endogenous mouse locus (Fig. S2A, Table S3). The levels of H3K4me2 were significantly increased at both the humanized and orthologous chimpanzee loci in each respective line compared to mouse (Fig. 2, S2, Table S3). As high levels of H3K4me2 coupled with low levels of H3K27ac are associated with weak enhancer activity (*23*), it is likely that the chimpanzee sequence is not acting as a strong enhancer in the limb bud overall, a finding further supported by the gene expression analyses described in Figure 3.

**Fig. 3.**
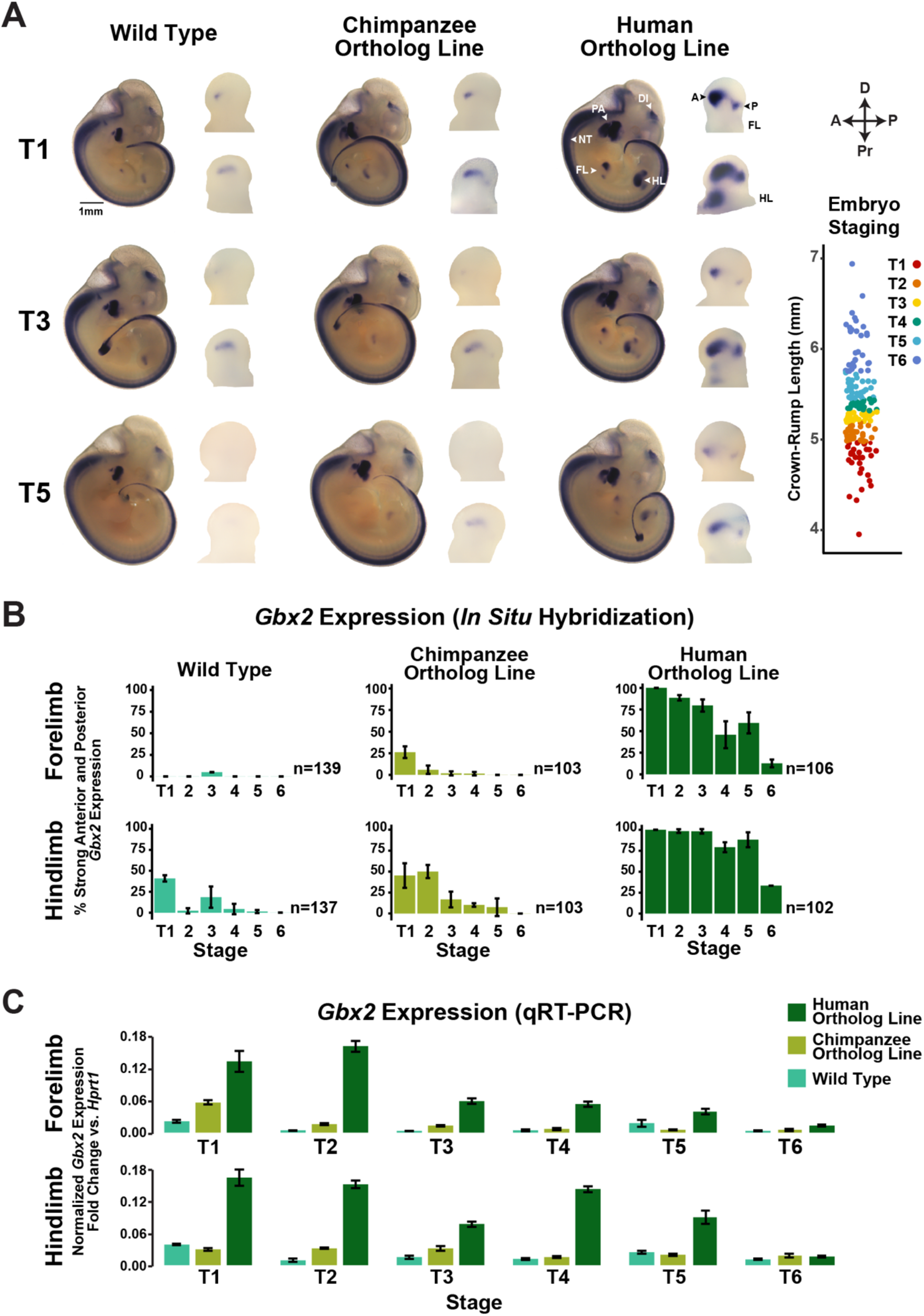
Spatial and temporal changes in *Gbx2* expression driven by *HACNS1* in humanized mouse embryos. **(A)** Spatial and temporal expression of *Gbx2* in *HACNS1* homozygous, chimpanzee ortholog line, and wild type E11-E12 embryos visualized by whole-mount *in situ* hybridization (ISH). Representative images are shown for each genotype at three fine-scale time points; see text and Fig. S3 for details on staging. Magnified views of *Gbx2* expression in limb buds are shown to the right of each embryo. Annotations of anatomical structures and developmental axes are indicated at the top right: FL = forelimb; HL = hindlimb; DI = diencephalon; NT = neural tube; PA = pharyngeal arch; A = anterior; P = posterior. The arrows at the top far right indicate the anterior-posterior (A-P) and proximal-distal (Pr-D) axes for the magnified limb buds. *Bottom right*: Crown-rump lengths for all embryos assayed for *Gbx2* mRNA expression by ISH. Each point indicates a single embryo. Colors denote each fine-scale time point (T1-T6). Photo Credit: Angeliki Louvi, Yale University. **(B)** Percentage of forelimbs or hindlimbs showing strong anterior and posterior expression of *Gbx2* at each time point in *HACNS1* homozygous, chimpanzee ortholog line, and wild type forelimb (top) and hindlimb (bottom). Error bars denote standard deviation for results obtained from three independent, blinded analyses; for scoring scheme with example images and full analysis see Fig. S3A (n = number of forelimbs or hindlimbs assessed). **(C)** Normalized *Gbx2* expression in pooled *HACNS1* homozygous, chimpanzee ortholog line, and wild type forelimb and hindlimb tissue across all six timepoints measured using qRT-PCR. Error bars denote standard deviation across three technical replicates.

Previous transgenic mouse enhancer assays also showed that *HACNS1* drives increased reporter gene activity in the pharyngeal arch compared to the chimpanzee ortholog (*10*). We therefore profiled H3K27ac and H3K4me2 in pharyngeal arch tissue from E11.5 embryos homozygous for either *HACNS1* or the chimpanzee ortholog and wild type embryos. We detected reproducible, significant enrichment of H3K27ac in the pharyngeal arch at the humanized and orthologous chimpanzee loci compared to input controls, but the H3K27ac signal at neither the human nor the chimpanzee ortholog locus was significantly different compared to the mouse endogenous locus (Fig. S2). We did, however, identify a significant gain of H3K4me2 signal in the pharyngeal arch at the humanized and orthologous chimpanzee loci compared to that at the mouse locus (Fig. S2, Table S3).

In order to identify downstream epigenetic changes resulting from *HACNS1* activation in *HACNS1* homozygous limb buds, we searched for other genome-wide gains of H3K27ac and H3K4me2 at enhancers and promoters. We identified a significant gain of H3K27ac in *HACNS1* versus wild type limb buds at the promoter of the nearby gene *Gbx2* (Fig. 2, S2, Table S3). While significant H3K27ac enrichment was found in all three lines at the *Gbx2* promoter compared to input controls, H3K27ac levels were not significantly increased at *Gbx2* in limb buds with the chimpanzee ortholog compared to wild type, indicating the gain of activity is specific to *HACNS1*. H3K4me2 was also enriched at the promoter of *Gbx2* in all three lines compared to input controls (Fig. S2D). After multiple testing correction, we did not identify any significant differentially marked regions outside of the *HACNS1-Gbx2* locus between either the humanized or chimpanzee ortholog line compared to wild type for each chromatin mark in either tissue (Table S3; Supplementary Materials). This may be due to a lack of statistical power to detect small differences in histone modification levels given the number of replicates in the analysis, or our use of whole limb tissues to map histone modification profiles, which could obscure spatially restricted changes.

*Gbx2* encodes a transcription factor with multiple functions during development. GBX2 has been implicated in midbrain and hindbrain development (*24, 25*), guidance of thalamocortical projections (*26, 27*), ear development (*28*), and pharyngeal arch patterning (*29*). *Gbx2* is expressed in developing mouse limb at E10.5; however, its role in limb development remains undetermined as no limb phenotype has been reported in *Gbx2* knockout mice (*25*). *HACNS1* and *GBX2* are located in the same topologically associated domain (TAD), and TADs have been shown to restrict enhancer interactions to genes within their boundaries (*30, 31*). The only significant increases in H3K27ac in the humanized limb detected in this TAD were at *HACNS1* and the *Gbx2* promoter (Table S3; Supplementary Materials). Together, these results suggest that *Gbx2* is a regulatory target of *HACNS1*, evoking the hypothesis that the gain of function in *HACNS1* might alter *Gbx2* expression in the humanized mouse limb.

### *HACNS1* drives spatial and quantitative changes in *Gbx2* expression in the limb bud

To visualize potential expression changes resulting from *HACNS1*-driven upregulation of the *Gbx2* promoter in humanized mouse embryos, we used *in situ* hybridization (ISH) (Fig.3). We analyzed *Gbx2* expression in >100 E11.5 embryos for each genotype (Fig. 3B and Fig. S3). In wild type embryos, we observed single foci of *Gbx2* expression in forelimb and hindlimb (Fig. 3A, left). In contrast, embryos homozygous for *HACNS1* showed substantially increased *Gbx2* expression in both forelimb and hindlimb (Fig. 3A, right). Embryos homozygous for the chimpanzee ortholog showed a weak increase in *Gbx2* expression compared to wild type (Fig. 3A and Figure S3A). *Gbx2* expression in *HACNS1* embryos was increased in two distinct anterior and posterior regions in the forelimb and hindlimb bud, as well as an anterior proximal region in the latter. Overall, *Gbx2* limb bud expression was temporally dynamic in embryos of all genotypes. Embryos from the same litter vary in developmental age such that individual embryos collected at E11.5 range from E11 to E12. Therefore, we established a fine staging scheme to characterize changes in *Gbx2* expression within this short developmental interval. We assigned embryos to 6 temporally ordered groups (designated T1-T6, and ranging from approximately 36 to 43 somites) according to crown-rump length and used a blinded approach to qualitatively assess staining patterns (Fig. 3A, Fig. S3A) (*32*).

We identified differences in the distribution of *Gbx2* expression in the forelimb and hindlimb buds of *HACNS1* embryos compared to both chimpanzee ortholog and wild type embryos across all 6 developmental time points (Fig. 3, S3). At the earliest time point (T1), we found that *Gbx2* was strongly expressed in distinct anterior-distal and posterior domains in *HACNS1* forelimb and hindlimb buds (Fig. 3, S3). Robust expression of *Gbx2* in *HACNS1* limb buds persisted through the remaining time points (up to T6), though the size of the anterior and posterior domains decreased over time. Strong expression of *Gbx2 in HACNS1* embryos persisted for a longer period of time in hindlimb than in forelimb, consistent with the delayed developmental maturation of the former (*33*). In addition, *HACNS1* embryos showed a hindlimb-specific anterior-proximal expression domain adjacent to the body wall across all 6 time points (Fig. S3B).

Compared to the robust expression in the humanized limb, *Gbx2* expression in limb buds from both the chimpanzee ortholog and wild type lines was weak and mostly evident at early time points (Fig. 3, S3A). Chimpanzee ortholog line embryos and wild type embryos both showed weak distal *Gbx2* expression foci in early forelimb and hindlimb that were generally restricted to the anterior portion of the limb bud (Fig. 3, S3A). Weak distal expression was primarily restricted to approximately T1-T2 in wild type forelimb but persisted until approximately T4 in a subset of embryos with the chimpanzee ortholog (Fig. 3, Fig S3A, top). Weak distal expression persisted in hindlimb through T5-T6 in both the chimpanzee line and wild type (Fig S3A, bottom). These findings suggest that the chimpanzee ortholog line exhibits a modest increase in *Gbx2* expression compared to wild type, potentially due to primate-rodent sequence differences affecting enhancer activity that our experimental design was intended to control for (see Supplemental Note). However, the *HACNS1* humanized line exhibits profound changes in *Gbx2* limb bud expression compared to both. Together, these findings suggest that *HACNS1* drives spatial and quantitative changes in *Gbx2* expression in the humanized limb, as well as a temporal extension of expression compared to wild type.

In addition to the forelimb and hindlimb bud, *Gbx2* was also expressed in the neural tube, diencephalon, and pharyngeal arch of embryos homozygous for *HACNS1* or the chimpanzee ortholog, and in wild type embryos (Fig. 3A, S3C) (*25, 27, 29*). Whereas *Gbx2* expression was primarily restricted to the first pharyngeal arch in embryos with the chimpanzee ortholog and in wild type embryos, we observed a dorsal expansion of *Gbx2* expression into the second pharyngeal arch in *HACNS1* embryos during T1-T5 (Fig. S3C). However, we chose to focus on limb due to a lack of comparative epigenetic profiling data in human pharyngeal arch and the absence of a significant gain in H3K27ac marking in *HACNS1* versus wild type pharyngeal arch.

In order to quantify the gain of *Gbx2* expression in humanized limbs, we used real-time quantitative reverse transcription PCR (qRT-PCR) in pooled forelimb and hindlimb buds from embryos homozygous for *HACNS1*, the chimpanzee ortholog, and the endogenous mouse locus at time points T1-T6. We found that *Gbx2* expression was increased in forelimb and hindlimb of *HACNS1* embryos versus both chimpanzee ortholog and wild type at all 6 time points (Fig. 3C). Although we detected an increase in *Gbx2* expression in forelimb and hindlimb of embryos with the chimpanzee ortholog versus wild type at early time points, this change was substantially weaker than that between *HACNS1* and wild type or *HACNS1* and the chimpanzee line. Consistent with our ISH results, *Gbx2* expression in humanized forelimb and hindlimb was strongest at the earliest time points and persisted longer in hindlimb than forelimb (Fig. 3C). While *Gbx2* expression declined over time in all three genotypes, it persisted longer in humanized forelimb and hindlimb (Tables S4, S5). To determine the significance of the effects of genotype and developmental age on *Gbx2* expression, we used analysis of variance (ANOVA). We determined that genotype, time point, and genotype-time point interaction effects were significant, indicating that *Gbx2* expression is significantly increased in humanized mouse (Tables S4, S5).

### *HACNS1* drives increased *Gbx2* expression in limb chondrogenic mesenchymal cells

In order to identify the specific cell types expressing *Gbx2* as well as genes that are potentially coregulated with *Gbx2* in the developing limb, we performed single-cell RNA-sequencing (scRNA-seq) in hindlimb cells from E11.5 embryos of all three genotypes. We focused on hindlimb for single-cell analysis as this tissue showed the most pronounced upregulation of *Gbx2* in spatial and quantitative expression analyses (Fig. 3, S3). Using the 10x Genomics scRNA-seq platform for cell barcoding, library preparation, and sequencing, we obtained transcriptomes from approximately 10,000 cells per genotype. We used the Seurat toolkit for data preprocessing and library size normalization (see Materials and Methods) (*34*). During pre-processing, we removed endothelial and blood cells (*Cd34*-positive; *Pf4*-positive; *Hbb*-positive), as our analysis indicated that these transcriptionally and developmentally distinct cell types do not express *Gbx2* in any of our datasets (*35, 36*). After normalization of the filtered data using the SCTransform method in Seurat and integration of data from all samples into a single dataset using the Seurat v3 integration workflow (see Materials and Methods), we performed clustering analysis on the integrated dataset to identify cell type categories present in all three genotypes (*34*). To visualize similarities between cells, we used Uniform Manifold Approximation and Projection for Dimensional Reduction (UMAP), a dimensionality reduction method for data visualization, followed by the Louvain method for community detection to identify cell subtypes (*37, 38*).

This analysis revealed three distinct groups: a) mesenchymal cell subtypes based on expression of the known markers *Sox9* (clusters 1, 3, 4), *Bmp4* (cluster 2)*, Shox2* (cluster 1), and *Hoxd13* (clusters 1-4) (Fig. 4A); b) non-mesenchymal cell types, including myogenic cells (cluster 5, *Myod*); and c) ectodermal cells (cluster 6, *Fgf8*) (Fig. 4A) (*39–44*). Furthermore, our analysis revealed finer separation of mesenchymal cells according to known limb patterning markers. We first examined the expression of known proximal-distal limb bud markers *Meis1, Hoxa11*, and *Hoxd13* (proximal, medial, and distal, respectively) (*45*). Cells expressing each of these markers showed a distinct localization in the UMAP embedding (*Meis1+* cells in the top left, *Hoxa11*+ cell in the center, and *Hoxd13*+ cells in the lower right; Fig. 4B, upper left), suggesting our analysis recovered transcriptional and cell-type transitions along the proximal-distal patterning axis (Fig. 4B).

**Fig. 4.**
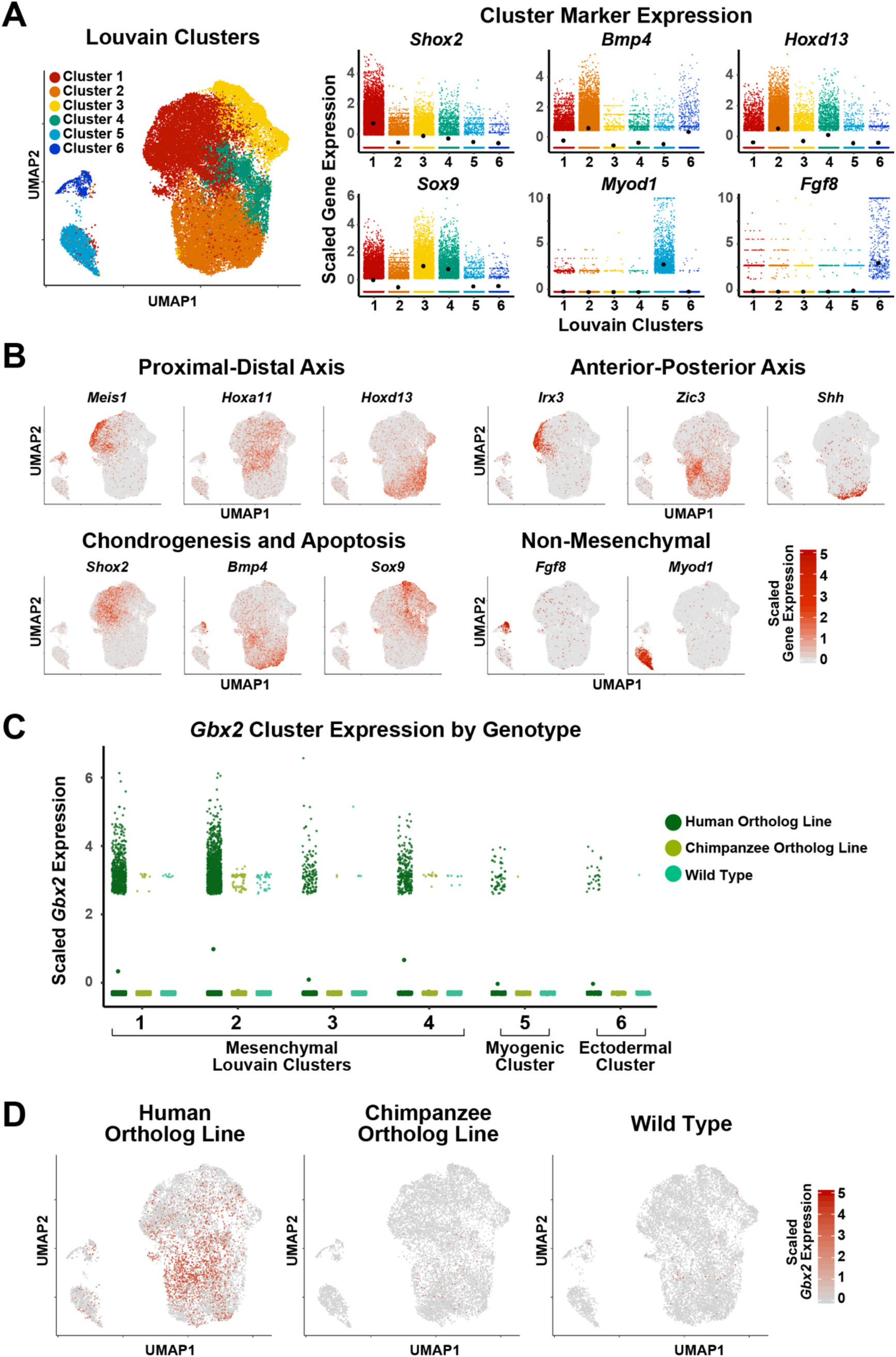
Single-cell transcriptome analysis of E11.5 hindlimb bud in *HACNS1* homozygous, chimpanzee ortholog line, and wild type embryos. **(A)** *Left:* UMAP embedding of *HACNS1* homozygous, chimpanzee ortholog line, and wild type cells. The colors indicate cell clusters identified by Louvain clustering. *Right:* Expression of known limb bud cell-type marker genes in each cluster. Black dots denote cluster mean expression. **(B)** UMAP embedding of hindlimb bud cells from *HACNS1* homozygous, chimpanzee ortholog line, and wild type, showing expression of proximal-distal, anterior-posterior, chondrogenesis-apoptosis, and non-mesenchymal markers. See text and Fig. S4 for details. **(C)** Expression of *Gbx2* in each Louvain cluster, separated by genotype. Dots denote cluster mean expression. **(D)** UMAP embeddings illustrating cells expressing *Gbx2* (indicated in red) in *HACNS1* homozygous, chimpanzee ortholog line, and wild type cells. All gene expression data shown in plots and UMAP embeddings (A-D) were imputed using ALRA and centered and scaled using z-scores (see Materials and Methods) (*75*).

We also found that the first axis of the UMAP embedding clearly recapitulated known gene expression gradients along the anterior-posterior limb bud axis based on expression of the anterior-proximal marker *Irx3,* the anterior marker *Zic3,* and the posterior-proximal marker *Shh* (Fig. 4B, upper right) (*39, 46–48*). Using markers of chondrogenic (*Sox9, Shox2*) versus non-chondrogenic (*Bmp4*) mesenchyme, we found that the second UMAP axis followed the chondrogenic versus interdigital apoptotic fate gradient (Fig. 4B) (*39, 41, 44*). We also found that the expression patterns of the aforementioned markers were broadly conserved between genotypes, with each genotype showing comparable subsets of proximal, distal, anterior, posterior, chondrogenic, and non-chondrogenic cell types (Fig. S4A, B). Collectively, our scRNA-seq analyses identified specific conserved cell types and spatial transcriptional gradients in the developing hindlimb bud across all three genotypes.

We then sought to define genotype-specific differences in *Gbx2* expression. In order to identify the cell types expressing *Gbx2* in humanized hindlimb bud, we examined the distribution of *Gbx2*-positive cells across cell clusters in all three genotypes. We found that *Gbx2* was upregulated in *HACNS1* hindlimbs versus chimpanzee ortholog and wild type hindlimbs, primarily in the mesenchymal cell clusters (clusters 1-4), consistent with the ISH and qRT-PCR expression analyses (Fig. 4C and Fig. 3). In *HACNS1* hindlimbs, 24% of cells expressed *Gbx2*, 96% of which were mesenchymal cells, whereas less than 1% of cells in chimpanzee ortholog and wildtype hindlimbs expressed *Gbx2* (see Materials and Methods; Table S6). *Gbx2* was primarily expressed in mesenchymal cells in both the chimpanzee ortholog line and wild type hindlimb, and most *Gbx2*-positive cells in each line were assigned to Louvain cluster 2. Only one non-mesenchymal cell from chimpanzee ortholog hindlimb and one non-mesenchymal cell from wild type hindlimb was *Gbx2-*positive (Fig. 4C; Table S6). UMAP embedding of cells revealed that *Gbx2-*positive cells in *HACNS1* hindlimb buds largely clustered within a distinct subset of mesenchymal cells belonging primarily to Louvain clusters 1, 2 and 4 (Fig. 4C, D; Table S6). The genotype-specific differences in *Gbx2* expression were consistent between the imputed and unimputed data as well as across individual replicates (Fig. S4C, D).

To identify genes whose expression is associated with *Gbx2*, we used k-Nearest-Neighbors Conditional-Density Resampled Estimate of Mutual Information (kNN-DREMI), which computes scores quantifying the strength of the relationship between two genes (*49, 50*). Using kNN-DREMI scores, we ranked each gene expressed in *HACNS1* humanized limb by the strength of its association with *Gbx2*. To determine if genes associated with *Gbx2* were enriched in particular functions, we then performed Gene Set Enrichment Analysis (GSEA) on this set of ranked genes. We found that *Gbx2* expression was associated with genes in several limb development-related ontologies, including “Cartilage Morphogenesis” (Kolmogorov-Smirnov (KS) P=1.61×10^−3^) and “Regulation of Chondrocyte Differentiation” (KS P=5.84×10^−3^); the latter overlapped considerably with “Collagen Fibril Organization” (KS P=1.30×10^−4^) (Fig. 5A, Table S7). These results indicate that in humanized mesenchyme, *Gbx2* is co-regulated with genes expressed in condensing mesenchymal cells destined to become chondrocytes (Fig. 5A).

**Fig. 5.**
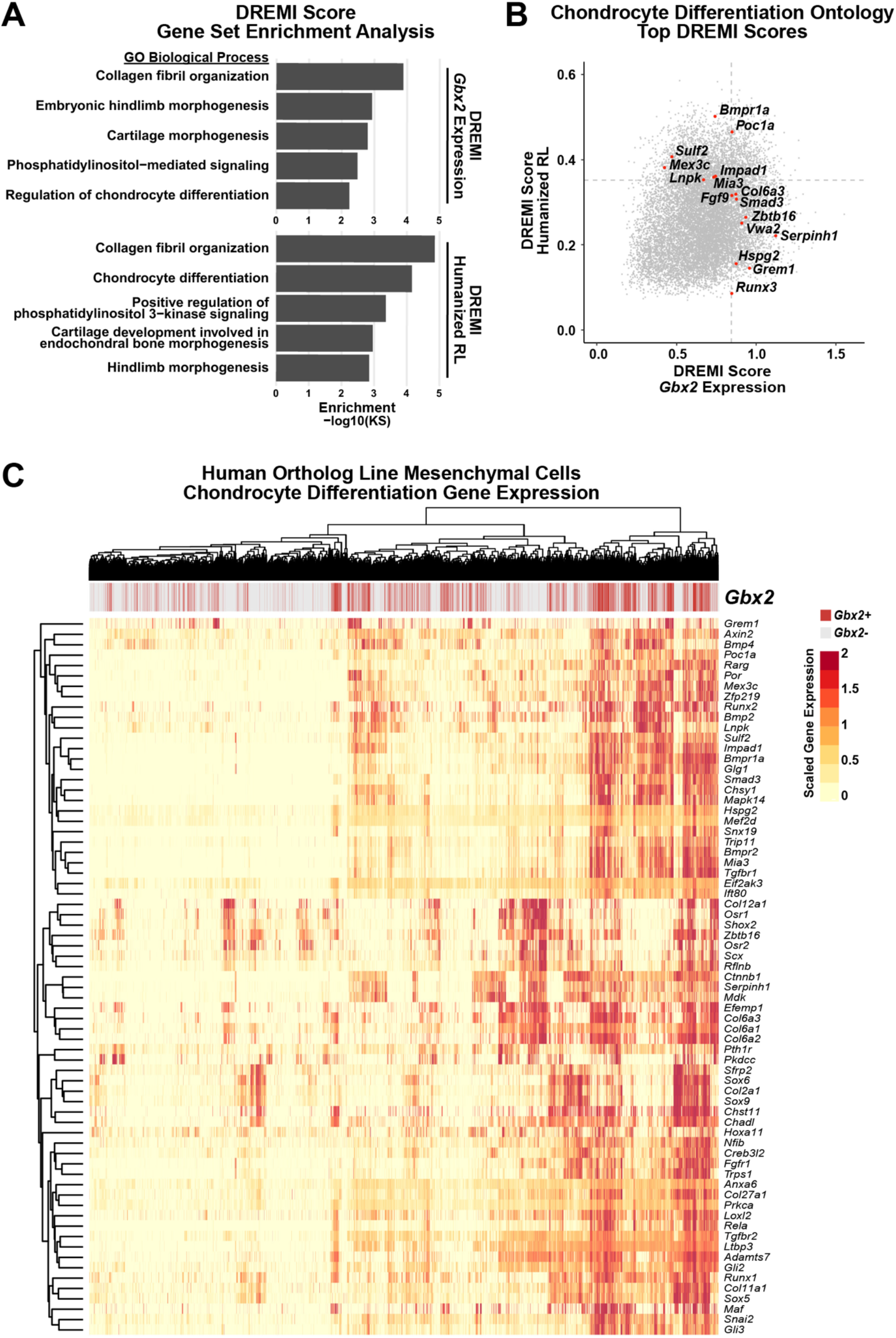
*Gbx2-*positive mesenchymal cell expression of chondrocyte differentiation markers in *HACNS1* homozygous limb bud. **(A)** Ontology enrichments of genes with expression associated with *Gbx2* expression (top) and the relative likelihood of the humanized condition (humanized RL, bottom) in *HACNS1* homozygous mesenchymal cells. The log-transformed Gene Set Enrichment Analysis Kolmogorov-Smirnov P value for each category is plotted on the x-axis. Ontologies shown are those overlapping in the *Gbx2* expression and humanized RL ontology enrichment lists. See also tables S7 and S8. **(B)** Humanized RL and *Gbx2* kNN-DREMI scores are plotted for all genes. Genes ranked in the top 20% of kNN-DREMI scores in the Chondrocyte Differentiation ontology (GO:0002062) for the union of the humanized RL and *Gbx2* kNN-DREMI analysis gene lists are colored in red and labeled. Dotted lines indicate the top 20% of values for each dataset. **(C)** Heatmap showing expression of genes belonging to the ontology “Chondrocyte Differentiation” (GO:0002062) in all *HACNS1* homozygous mesenchymal cells (Louvain clusters 1-4). Hierarchical clustering was used to determine the order of cells (in columns) and genes (in rows). The bar at the top of the heatmap shows *Gbx2*-positive and *Gbx2*-negative cells in red and gray, respectively. The gene expression values shown are imputed using ALRA and centered and scaled using z-scores (see Materials and Methods) (*75*).

We also used Manifold Enhancement of Latent Dimensions (MELD) to quantify the differences between humanized and non-humanized limb bud transcriptional profiles. MELD is an unsupervised learning algorithm and is therefore an orthogonal approach that is naïve to our identification of *Gbx2* as the target of *HACNS1* (*51*). MELD uses graph signal processing to quantify the relative likelihood of observing each cell in each of multiple experimental conditions based on its transcriptional profile. In this case, MELD is used to quantify the relative likelihood (RL) of observing a cell in the *HACNS1* hindlimb cells versus the chimpanzee ortholog or wild type hindlimb. Rather than explicitly classifying genes as differentially expressed in one condition versus another, the RL value can be used to identify trends in gene expression across cells that are associated with the humanized condition (*51*).

To identify overall gene expression patterns characteristic of humanized hindlimb bud cells, we used kNN-DREMI to associate gene expression with the humanized condition (humanized RL) calculated by MELD. We then used the resulting gene rankings to identify enriched biological functions via GSEA, as described above for *Gbx2* expression. We found that genes associated with both humanized RL and *Gbx2* expression converged on related biological processes. Performing GSEA using genes ranked by mutual information with humanized RL revealed significant enrichment of the “Chondrocyte Differentiation” ontology (KS P=7.00×10^−5^), along with four other categories also significantly enriched in the *Gbx2* expression analysis: “Hindlimb Morphogenesis” (KS P=1.42×10^−3^), “Cartilage Development Involved in Endochondral Bone Morphogenesis” (KS P=1.11×10^−3^), “Collagen Fibril Organization” (KS P=1.40×10^−5^), and “Positive Regulation of Phosphatidylinositol 3-kinase Signaling” (KS P=4.40×10^−5^), of which the last two are also implicated in chondrocyte differentiation (Fig. 5A, Table S8) (*52, 53*). “Collagen Fibril Organization” is the most significantly enriched GO term for genes associated with humanized RL and is the second most enriched for genes associated with *Gbx2* expression. The top GO term for genes associated with *Gbx2* expression, “Roof of Mouth Development,” (KS P=1.10×10^−5^), shares >20% of its genes with “Embryonic Hindlimb Morphogenesis” (KS P=1.18×10^−3^) (*54*). This illustrates that many genes involved in limb development are also implicated in craniofacial development, and likely accounts for why craniofacial development-related GO terms were also enriched in our analysis.

These findings led us to examine the expression patterns of chondrocyte differentiation-related genes in humanized mesenchymal cells belonging to Louvain clusters 1-4 (Fig. 5C). We clustered *HACNS1* mesenchymal cells by humanized RL and *Gbx2* expression and examined the expression of the “Chondrocyte Differentiation” ontology genes within *Gbx2*-positive cells (Fig. 5C). This clustering analysis revealed higher expression of positive regulators of chondrocyte differentiation (e.g. *Sox9*, *Col2a1*, *Bmp2*, and *Runx2)* specifically in *Gbx2*-positive versus *Gbx2*-negative humanized mesenchymal cells, supporting that *HACNS1-*driven upregulation of *Gbx2* occurs in chondrogenic cells destined for digit formation (Fig. 5C) (*53, 55–57*). We also identified a subset of *Gbx2*-positive humanized cells that were also positive for *Bmp4*, which is expressed in the apoptotic interdigital domains (*58*). These findings suggest that upregulation of *Gbx2* impacts the interdependent pathways of digit condensation and interdigital cell fate specification required for digit morphogenesis.

### Morphometric analysis of humanized mouse limbs

To determine if *Gbx2* upregulation and downstream transcriptional changes in *HACNS1* limb buds affect digit formation or overall limb morphology, we performed morphometric analysis of skeletal preparations for embryos homozygous for *HACNS1*, embryos homozygous for the chimpanzee ortholog, and wild type embryos (Materials and Methods). We performed morphometric analysis at E18.5 in order to capture any major phenotypic effects of humanization that occurred by the end of embryonic skeletogenesis (*20*). We did not detect gross morphological differences among genotypes; the three major limb segments (autopod, zeugopod, and stylopod) were present in both *HACNS1* skeletons and chimpanzee ortholog skeletons (Fig. S6). We also examined digit length (normalized to body size based on the length of the ossified humerus), and intradigital (phalange to metacarpal or metatarsal length) and interdigital ratios. Again, we found no significant differences in digit length or autopod proportions between genotypes (Fig. S6, Tables S9-S11).

## Discussion

Understanding how uniquely human genetic changes altered developmental processes is essential to understanding the evolution of our species. Here we investigated the role of the Human Accelerated Region *HACNS1* in human limb evolution by directly interrogating its biological functions in a humanized mouse model. This *in vivo* approach enabled us to identify spatial and temporal changes in gene expression driven by *HACNS1* and to characterize the specific cell types affected by these changes, providing insight into the developmental processes modified due to human-specific alterations in enhancer activity.

First, we determined that *HACNS1* is active in the mouse genomic context, recapitulating the significant level of H3K27ac marking previously observed in the developing human limb in the *trans-*regulatory environment of the developing mouse limb. Second, we found that *Gbx2* exhibited increased promoter activity in the humanized limb bud, strongly supporting that it is regulated by *HACNS1*, and then demonstrated that *HACNS1* drives robust changes in *Gbx2* expression in the forelimb and hindlimb bud. These findings support the long-standing hypothesis that discrete regulatory changes altering expression of pleiotropic developmental regulators in specific tissues contribute to the evolution of phenotypic differences – in the case of this study, molecular phenotypes - across species (*18, 19*). Third, by performing scRNA-sequencing, we identified the spectrum of distinct and specific cell types that show upregulation of *Gbx2* in the developing limb, as defined by transcriptome signatures. This analysis not only established that *Gbx2* is expressed in mesenchymal cells in the limb bud, but also placed these cells in the developmental process of chondrogenesis. By both characterizing *Gbx2*-positive humanized cells and identifying overall expression trends associated with the humanized condition without reference to *Gbx2* expression, we implicated changes in *Gbx2* regulation in chondrocyte differentiation, a critical process in digit formation.

We found that the human-specific gain of function in *HACNS1* drives quantitative, spatial, and temporal changes in *Gbx2* expression in the humanized mouse limb bud. One hypothesis consistent with these findings is that *HACNS1* acts to both spatially expand and prolong *Gbx2* expression and its potential effects on chondrocyte differentiation in the developing limb. *Gbx2* is expressed in wild type mouse limb at E10.5, and the mouse ortholog of *HACNS1* is marked by H3K27ac at this time point, suggesting it is contributing to *Gbx2* regulation (*25, 59, 60*). However, the mouse ortholog is annotated as a poised enhancer in the E11 limb bud, consistent with the weak expression of *Gbx2* we observe in E11-E12 wild type limb buds (*61*). In contrast, *HACNS1* shows robust H3K27ac marking in E11.5 limb, and *Gbx2* is strongly expressed in the E11 humanized limb bud. These results, coupled with our finding that *Gbx2* is primarily expressed in limb mesenchymal cells in all three genetic backgrounds we interrogated, suggests that the gain of function in *HACNS1* may be modifying an ancestral *Gbx2* regulatory program in the limb. Further insight into this question will require deciphering the *Gbx2* regulatory network in both humanized and wild type limbs, including identifying downstream targets of *Gbx2* and upstream regulators of *HACNS1* and the orthologous mouse enhancer. We note that several of the transcription factors whose predicted binding sites are only present in *HACNS1* have been implicated in limb bud patterning and chondrogenesis at E11, including *Ets1* and *Gabpa* (*62*), *Tfap2B* (*63*), and *Runx2* (Table S2) (*64*).

The modest, but observable, differences in *Gbx2* expression in the chimpanzee ortholog line compared to wild type may be due to primate-rodent sequence changes that altered ancestral enhancer activity at the *HACNS1* locus, predating the human-specific changes we focused on in this study. Although *HACNS1* itself shows very high levels of sequence conservation with the chimpanzee and mouse orthologs, the flanking human and chimpanzee sequences we included in our knock-in models harbor multiple differences, including single nucleotide substitutions and gaps, relative to the endogenous mouse locus (Fig. 1B; Supplemental Note). These sequence changes may have contributed either to gain of function at the *HACNS1* locus during primate evolution, or to loss of function on the rodent lineage. Distinguishing between these mechanisms will require *in vivo* genetic studies of *HACNS1* orthologs from multiple primate, rodent and outgroup species, and of primate and rodent ancestral orthologs inferred by ancestral sequence reconstruction.

Our morphological studies in the *HACNS1* mouse model also have several limitations that we note here. We did not identify major changes in skeletal morphology at E18.5 that were associated with humanization. However, this does not preclude subtle changes in limb length or other features in adult mice, or changes in soft tissues that would not be detected using skeletal preparations. We also did not exhaustively characterize other tissues, including the pharyngeal arch and diencephalon, in which *Gbx2* has known developmental functions (*24–27, 29*). Although we did not observe overt craniofacial phenotypes in *HACNS1* humanized mice, the dorsal expansion of *Gbx2* expression into the second pharyngeal arch that we detected in this background may result in more subtle developmental effects that could be explored in future studies.

We have shown that humanized mouse models represent a viable and fruitful approach for studying gene regulatory mechanisms relevant for human evolution within the complete genomic, tissue-level, and developmental framework of a living organism. That the molecular phenotypes we observed did not produce an overt morphological phenotype is not entirely surprising. Genetic changes in any one enhancer are unlikely to be sufficient to replicate human-specific morphological changes entirely in an experimental model. The evolution of uniquely human physical traits likely entailed modifications in the expression of many genes, potentially driven by multiple HARs and other human-specific genetic changes. Our study provides insight into how a single HAR alters gene regulation and expression at critical developmental time points, yielding an important entry point for understanding the larger developmental networks that changed during human limb evolution, of which *Gbx2* is a part. Humanized mouse models offer the means to study additional HARs contributing to human-specific phenotypes at once, either through intercrossing mouse lines harboring edited unlinked loci or by iterative editing of one locus, allowing us to expand our understanding of human limb evolution. Our study thus establishes a framework for using humanized mouse models to link individual sequences that arose on the human lineage since species divergence to the unique traits that distinguish our species.

## Materials and Methods

### Mouse Line Generation and Validation

The *HACNS1* and chimpanzee ortholog lines were generated at the Yale Genome Editing Center using standard techniques in modified ES cells (*65*). C57BL/6J-*A^w-J^*/J mouse ES cells were edited by electroporation of a GFP cloning vector containing human (1,241 bp) or chimpanzee (1,240 bp) sequence flanked by C57BL/6J mouse sequence homology arms, floxed pPGKneo vector, and diphtheria toxin sequence (Fig. S1A) (*66*). The genomic coordinates of the human (hg19) and mouse (mm9) sequences used in the editing constructs, including the mouse homology arm sequences, are listed in Table S1 (*67*). G0 chimeras were backcrossed to wild type C57BL/6J (RRID: IMSR_JAX:000664) and crossed with an actin-Cre C57BL/6J mouse line to remove the neo cassette. All mice used in our analysis were from F9 or later generations. All animal work was performed in accordance with approved Yale IACUC protocols.

Genotyping primers specific to *HACNS1*, chimpanzee, and mouse orthologs are listed in Table S12. Cloning primers listed in Table S12 were used to amplify edited loci for cloning and Sanger sequencing for comparison to the hg19 or panTro4 sequence. Sanger sequencing data is available at noonan.ycga.yale.edu/noonan_public/Dutrow_HACNS1/. The sequence identity between the human (hg19, chr2:236773456-236774696) and chimpanzee alleles (panTro4, chr2B:241105291-241106530) is 98.2% (22 substitutions total, of which 15 are fixed in humans). Human-specific substitutions were defined as fixed if the derived allele frequency in dbSNP (v153) was >= 0.9999 and if the ancestral sequence state was conserved between chimpanzee, rhesus macaque, orangutan, and marmoset. We provide a detailed analysis of sequence differences between the human, chimpanzee and mouse orthologs in the Supplemental Note (Supplementary Materials). *HACNS1-GBX2* locus TAD coordinates (hg19 chr2:236655261-237135261) are from H1 human ES cell Hi-C data; *HACNS1* and *GBX2* are present in the same TAD and *GBX2* is the only annotated protein-coding gene whose promoter is included in this TAD (*30*).

Copy number verification qPCR was performed using genomic DNA from three F9 mice from each line using Power SYBR Green Mastermix (Thermo Fisher Scientific #4368577) and the StepOnePlus Real-Time PCR System (Applied Biosystems) with primers listed in Table S12. All biological replicates of each genotype were run in triplicate and Ct values of each were normalized to a control region on a different chromosome (see Table S12).

### Chromatin Immunoprecipitation, ChIP-qPCR and ChIP-seq

Tissue for chromatin preparation was collected from E11.5 forelimb and hindlimb bud pairs or pharyngeal arch tissue from *HACNS1* and chimpanzee ortholog line heterozygous crosses to obtain pooled, litter matched limb bud or pharyngeal arch samples for all three genotypes (*HACNS1* homozygous, chimpanzee ortholog line, and wild type). Two biological replicates were used per genotype per tissue, each with tissue pooled from three embryos. Pooled tissue was crosslinked and sonicated as previously described (*68*). Chromatin for each genotype, tissue, and replicate was used for H3K27ac or H3K4me2 immunoprecipitation using Active Motif #39133 (RRID: AB_2561016) and Active Motif #39913 (RRID: AB_2614976) antibodies as previously described (*68, 69*). ChIP-qPCR was performed using Power SYBR Green Mastermix (Thermo Fisher Scientific #4368577) with primers listed in Table S13. Samples were sequenced (2×100 bp) using standard Illumina protocols on an Illumina HiSeq 4000 (RRID: SCR_016386). To control for batch effects, all samples of the same tissue type were multiplexed and sequenced on a single lane.

Reference genomes edited to replace the mouse ortholog of *HACNS1* with the human or chimpanzee sequence were built using Bowtie (v2.2.8; RRID: SCR_005476) (*70*). ChIP-seq raw reads were aligned to the mm9, mm9 with chimpanzee ortholog, or humanized mm9 reference genome using Bowtie with --sensitive and --no-unal settings (*71*). GC composition was assessed using fastQC and showed that GC content and bias were consistent across all experiments (*72*). Tag directories for each experiment were generated using makeTagDirectory in HOMER with default settings and standard normalization to 10 million tags, and were used to generate bigwig files for visualization with makeUCSCfile (*21*). All peaks were called with HOMER, and all differential peaks were called with DESeq2 implemented in HOMER (see Extended Methods for details) (*21, 22*). The complete datasets of all peaks tested in differential analyses can be found at noonan.ycga.yale.edu/noonan_public/Dutrow_HACNS1/.

### RNA Extraction and qRT-PCR

E11-E12 embryos were collected from six *HACNS1* homozygous, chimpanzee ortholog line, or wild type litters generated by crossing homozygous animals for each line. All embryos within each genotype group were ordered based on stage and were divided into six timepoint groups per genotype consisting of forelimb or hindlimb buds from 4-6 pooled embryos per time point per genotype per tissue. RNA was purified using the Qiagen miRNeasy Kit (#74106). Invitrogen Superscript III Reverse Transcription Kit (#18080-051) was used to prepare cDNA from each sample. qPCR with the resulting cDNA was performed using Power SYBR Green Mastermix (Thermo Fisher Scientific #4368577). All samples were analyzed in triplicate using primers listed in Table S14 and Ct values of *Gbx2* were normalized to *Hprt1*.

### Whole mount *In Situ* Hybridization

E11-E12 mouse embryos were collected from *HACNS1* homozygous (n=7 litters), chimpanzee ortholog line (n=8 litters), and wild type (n=12 litters) homozygous crosses. Embryos were fixed and hybridized with the same preparation of antisense *Gbx2* mRNA probe under identical conditions as previously described (*71, 73*). The *Gbx2* probe used for hybridization contains the full mouse consensus CDS sequence (CCDS15150.1; NCBI CCDS Release 23). The 178 embryos (55 from the humanized line, 52 from the chimpanzee ortholog line, and 71 from wild type) were divided into sextiles based on crown-rump length and assessed for staining pattern by three individuals blinded to genotype under a stereo microscope (Leica S6D). See Extended Methods for further details regarding the scoring scheme used for qualitative assessment of expression and Fig. S3A for example images of staining patterns. Representative images were taken using a Zeiss Stemi stereomicroscope. Images and associated data are available at noonan.ycga.yale.edu/noonan_public/Dutrow_HACNS1/.

### Single-Cell RNA-Sequencing

#### Sample Preparation

Tissue for scRNA-seq was collected at E11.5 from two human ortholog line homozygous litters, two chimpanzee ortholog line homozygous litters, and two wild type litters. Embryos were staged as previously described in order to obtain samples from stage-matched T3 embryos from each genotype. Left hindlimb buds from three embryos per genotype per replicate were pooled. Following dissection, cells were dissociated and processed for library preparation using Chromium Single Cell 3ʹ GEM, Library & Gel Bead Kit v3 (10X Genomics PN-1000075) (see Extended Methods for details). Libraries were sequenced (2×75 bp) on an Illumina HiSeq 4000 (RRID: SCR_016386). To control for batch effects, all samples were multiplexed across all lanes. Count matrices were produced from raw sequencing data using the Cell Ranger v3.0.2 package from 10X Genomics (RRID: SCR_017344).

#### Data Pre-processing, Normalization, and Intergration

Matrices from the 10x Cell Ranger platform were filtered and preprocessed using Seurat v3.0.1 (RRID: SCR_016341) (*34*). For pre-processing, cells were clustered to remove low quality, blood, and endothelial cells (see Extended Methods) (*34, 38*). All subsequent normalization and integration steps after pre-processing were performed with raw counts for all cells retained after pre-processing (see Table S15). Cell cycle scores, percent mitochondrial gene expression and nUMI values were regressed using SCTransform and all normalized datasets containing all genes from individual samples were integrated using Seurat (*34, 74*). PCA, UMAP, and Louvain clustering were implemented in Seurat (*34, 37*). Normalized data from all samples combined were used for imputation using ALRA (*75*). For detailed description of data processing see Extended Methods.

#### MELD, MAGIC, kNN-DREMI Analyses

Cells belonging to mesenchymal cell clusters (clusters 1-4, see Fig. 4A, C) from all genotypes were used for MELD, MAGIC, kNN-DREMI, and Gene Set Enrichment Analysis (GSEA). MELD was run on one-hot vectors for each genotype independently using default parameters (*51*). MAGIC was performed using the same graph signal as MELD (*50*). kNN-DREMI was used to identify genes with expression levels associated with either *Gbx2* expression in humanized hindlimb or cells with increased humanized RL as calculated using MELD (*49*). See Extended Methods for further details.

#### Gene Set Enrichment Analysis

GSEA was performed using topGO v.2.34.0 (RRID: SCR_014798) on all expressed genes that were ranked by *Gbx2-*DREMI or humanized RL-DREMI score from the aforementioned humanized mesenchymal cell kNN-DREMI analysis (*76*). Significant nodes were identified using the Kolmogorov–Smirnov test and *elim* algorithm. Ontologies listed in Tables S7 and S8 are the top 30 nodes with fewer than 100 annotated genes (to remove non-specific categories) and at least one gene in the top 20% of DREMI scores. Heatmap hierarchical clustering was performed using pheatmap v1.0.12 (RRID: SCR_016418) (*77*).

### Skeletal Staining

E18.5 skeletons from two litters from each of *HACNS1* homozygous, chimpanzee ortholog line, and wild type homozygous crosses (n=48 embryos) were stained with Alcian Blue and Alizarin Red as previously described (*65*). Skeletons were imaged under a stereo microscope (Leica S6D) and measured blinded to genotype using ImageJ 2.0.0 (for complete list of measurements taken see Extended Methods). Digit length was calculated as the sum of all metacarpal/metatarsal and phalanx segments. Raw measurements and digit length were normalized to the length of ossified humerus or femur for forelimb or hindlimb digits, respectively. Phalange to metacarpal ratio was calculated as the ratio of the sum of the phalange lengths of each digit to the corresponding metacarpal/metatarsal segment. Interdigital ratios were calculated using raw digit lengths. Raw measurements and images are available at noonan.ycga.yale.edu/noonan_public/Dutrow_HACNS1/.

### ANOVA Analysis for Gene Expression and Morphometry

ANOVA analysis was performed with the lme4 package in R (RRID: SCR_015654) using default parameters to dissect the effects of genotype on *Gbx2* expression (qRT-PCR data) and limb segment length (morphometric data) (*78*). Comparisons adjustments were performed using the Benjamini & Hochberg method (*79*). See Extended Methods for details regarding specific analyses.

## Supporting information

Dutrow et al 2021 Supplemental Tables

## Acknowledgments

**Funding** This work was supported by a grant from the National Institute of General Medical Sciences (NIGMS) (R01 GM094780, to J.P.N.), funds from the Yale School of Medicine (to J.P.N.), and funds from the Yale Program on Neurogenetics (to A.L.). E.V.D. was supported in part by NIGMS training grant T32 GM007499. D.E. was supported by an NIH F32 Postdoctoral Fellowship (NIGMS) (F32 GM106628). A.K. was supported by an NSF Graduate Research Fellowship (DGE-1122492). K.Y. was supported by an NSF Graduate Research Fellowship (DGE-1752134). S.U. was supported by a Research Fellowship (352711928) from the Deutsche Forschungsgemeinschaft (DFG). M.K. was supported by a Research Fellowship (387495052) from the Deutsche Forschungsgemeinschaft (DFG). D.B. was supported by an NIH F31 Predoctoral Fellowship (NICHD) (F31 HD097958). S.K. was supported by NIH grants R01 GM135929 and R01 GM130847, and Chan-Zuckerberg Initiative grants 182702 and CZF2019-002440. This research program and related results were also made possible by the support of the NOMIS foundation (to J.P.N.).

**Author Contributions** E.V.D. and J.P.N. conceived of and designed the study with input from D.E.; E.V.D. performed the mouse line validation, chromatin immunoprecipitation, qRT-PCR, single cell RNA-sequencing, and skeletal staining experiments; E.V.D. and A.L. carried out the *in situ* hybridization experiments with assistance from A.A.K. and M.K.; D.E. and T.N. initially generated the mouse lines used in the study. Data analysis and interpretation for the single cell RNA-sequencing experiment was performed by E.V.D. with input from K.Y., D.B.B. and S.K. S.U. performed the ANOVA analyses. E.V.D., A.L. and J.P.N. wrote the manuscript with input from all authors.

**Data availability** The Gene Expression Omnibus accession number for the data reported in this paper is GSE141471. Access to all additional data needed to evaluate the conclusions in the paper is provided in the paper and the Supplementary Materials.

## Supplementary Materials for

**This file includes:**

Extended Methods

Supplemental Note

Figs. S1 to S6

**Other Supplementary Materials for this manuscript include the following:**

Table S1. Genomic Sequence Coordinates of Editing Construct Template DNA

Table S2. Predicted Transcription Factor Binding Site Changes in *HACNS1*

Table S3. ChIP-seq Significant Differential Peaks

Table S4. Time Series qRT-PCR Forelimb ANOVA Analysis

Table S5. Time Series qRT-PCR Hindlimb ANOVA Analysis

Table S6. scRNA-seq *Gbx2* Expression Summary

Table S7. *Gbx2* kNN-DREMI GSEA Results

Table S8. Humanized RL kNN-DREMI GSEA Results

Table S9. Normalized Digit Length ANOVA

Table S10. Phalange to Metacarpal/Metatarsal Ratio ANOVA

Table S11. Interdigital Ratio ANOVA

Table S12. Oligonucleotides for Genotyping, Cloning, and Copy Number Analysis

Table S13. Oligonucleotides Used for ChIP-qPCR

Table S14. Oligonucleotides Used for qRT-PCR

Table S15. scRNA-seq Sample Summary

### Extended Methods

#### Chromatin Immunoprecipitation, ChIP-qPCR and ChIP-seq

Reference genomes edited to replace the mouse ortholog of *HACNS1* with the human or chimpanzee sequence were built using Bowtie (v2.2.8; RRID: SCR_005476) (*70*). ChIP-seq raw reads were aligned to the mm9, mm9 with chimpanzee ortholog, or humanized mm9 reference genome using Bowtie with --sensitive and --no-unal settings (*71*). GC composition was assessed using fastQC and showed that GC content and bias were consistent across all experiments (*72*). Tag directories for each experiment were generated using makeTagDirectory in HOMER with default settings and standard normalization to 10 million tags, and were used to generate bigwig files for visualization with makeUCSCfile (*21*). All peaks were called with HOMER (v4.9.1 RRID: SCR_010881) using default settings for --histone (IP vs input fold change=4, p=0.0001, peak size=500, minDist=1000) (*21*). All differential peaks were called with DESeq2 implemented in HOMER using getDifferentialPeaksReplicates.pl with default settings (fold change cutoff =2, FDR cutoff = 5%); briefly, reads from each comparison are pooled, with ChIP and inputs pooled separately, such that new peaks are called and used for quantitative comparison between genotypes (*21, 22*). The complete datasets of all peaks tested in differential analyses can be found at noonan.ycga.yale.edu/noonan_public/Dutrow_HACNS1/.

#### Whole mount *In Situ* Hybridization

E11-E12 mouse embryos were collected from *HACNS1* homozygous (n=7 litters), chimpanzee ortholog line (n=8 litters), and wild type (n=12 litters) homozygous crosses. Embryos were fixed and hybridized with the same preparation of antisense *Gbx2* mRNA probe under identical conditions as previously described (*71, 73*). The *Gbx2* probe used for hybridization contains the full mouse consensus CDS sequence (CCDS15150.1; NCBI CCDS Release 23). The 178 embryos (55 from the humanized line, 52 from the chimpanzee ortholog line, and 71 from wild type) were divided into sextiles based on crown-rump length and assessed for staining pattern by three individuals blinded to genotype under a stereo microscope (Leica S6D). See Fig. S3A for example images of staining patterns representing the scoring scheme used for qualitative assessment of expression. Embryos were assessed for one of eleven categories of *Gbx2* expression pattern: 1: anterior and posterior (AP); 2: anterior distal and posterior distal (APD); 3: distal (D); 4: anterior distal (AD); 5: anterior (A); 6: weak anterior and posterior (APL); 7: weak anterior (AL); 8: weak distal (DL); 9: weak anterior and posterior distal (APDL); 10: weak anterior distal (ADL); 11: no staining (N). Categories were merged for clarity in Fig. S3A in the following manner: categories 1-3: anterior and posterior; categories 4-5: anterior only; categories 6-10: weak staining.

Representative images were taken using a Zeiss Stemi stereomicroscope. Images and associated data are available at noonan.ycga.yale.edu/noonan_public/Dutrow_HACNS1/.

#### Single-Cell RNA-Sequencing

##### Sample Preparation

Tissue for scRNA-seq was collected at E11.5 from two human ortholog line homozygous litters, two chimpanzee ortholog line homozygous litters, and two wild type litters. Embryos were staged as previously described in order to obtain samples from stage-matched T3 embryos from each genotype. Left hindlimb buds from three embryos per genotype per replicate were pooled. Following dissection, the tissue was immediately placed in CMFSG saline–glucose solution (1× Calcium–magnesium-free phosphate buffered saline from Thermo Fisher Scientific #21-040-CV with 0.1% glucose from Corning 45% Glucose #45001-116) on ice. Gibco TrypLE Express digestion solution was used for cellular dissociation (Thermo Fisher Scientific # 2605010). The dissociation reaction was stopped using 1xDMEM (ATCC 30-2002) with 10% heat-inactivated Fetal Bovine Serum (Sigma-Aldrich #F4135). The dissociated cells were filtered through a 40 μM strainer and harvested by centrifugation at 4**°**C. Cells were washed and resuspended in 1x Calcium–magnesium-free phosphate buffered saline (Thermo Fisher Scientific #21-040-CV) with 0.04% BSA (Sigma-Aldrich #SRE0036). Cell number and viability were estimated on a Countess II Automated Cell Counter prior to library preparation of 10,000 cells (estimated cell recovery from 16,000 input cells) per sample using Chromium Single Cell 3ʹ GEM, Library & Gel Bead Kit v3 (10X Genomics PN-1000075). Libraries were sequenced (2×75 bp) on an Illumina HiSeq 4000 (RRID: SCR_016386). To control for batch effects, all samples were multiplexed across all lanes. Count matrices were produced from raw sequencing data using the Cell Ranger v3.0.2 package from 10X Genomics (RRID: SCR_017344).

##### Filtering and Preprocessing

Matrices from the 10x Cell Ranger platform were filtered and preprocessed using Seurat v3.0.1 (RRID: SCR_016341) (*34*). Prior to the generation of Seurat objects, *Xist* gene counts were eliminated in order to avoid clustering by sex within mixed sample populations. Genes expressed in fewer than 5 cells per sample were removed. Cells with greater than 7.5% or 2% counts from mitochondrial genes or hemoglobin genes, respectively, were removed. Cells with total gene count (nGene) z-scores less than −1 (corresponding to approximately 700 or fewer detected genes) or greater than 4 (corresponding to approximately 6000 or greater detected genes) were removed, as were cells with total UMI count (nUMI) z-scores greater than 7 (corresponding to approximately 50,000 or greater detected UMIs; see Fig. S5). One chimpanzee ortholog line replicate was removed during pre-processing due to high overall mitochondrial gene expression, indicative of low viability. Prior to data integration, expression values from each sample were normalized based on library size for pre-processing purposes only using the Seurat tool NormalizeData (*34*). Louvain clustering as implemented in Seurat was performed for pre-processing purposes only using FindVariableFeatures, ScaleData, RunPCA, FindNeighbors, and FindClusters in order to remove endothelial cell clusters (*Cd34*-positive and *Pf4*-positive), clusters characterized by aberrant mitochondrial gene expression (low *mt-Co1*), and transcriptionally distinct clusters containing fewer than 30 cells per sample (*34, 38*). The numbers of cells remaining after pre-processing for each sample are listed in Table S15.

##### Data Normalization and Integration

All subsequent normalization and integration steps after pre-processing were performed with raw counts for all cells retained after pre-processing (see Table S15). Cell cycle scores were computed using CellCycleScoring in Seurat to regress out the difference between G2M and S phases, effectively preserving differences between cycling and non-cycling cells while reducing differences related to cell cycle amongst proliferating cells (*34*). In addition to cell cycle scores, percent mitochondrial gene expression and nUMI values were regressed using SCTransform (SCT) in order to reduce the effects of sequencing depth and minor differences in mitochondrial DNA expression related to viability (*74*). All SCT normalized datasets containing all genes from individual samples were integrated using SelectIntegrationFeatures, PrepSCTIntegration, FindIntegrationAnchors, and IntegrateData (*34, 74*).

Following integration, the combined dataset was randomly down-sampled to contain a maximum of 10,000 cells per genotype prior to embedding and clustering using SubsetData in Seurat (*34*). PCA, UMAP, and Louvain clustering were implemented in Seurat using RunPCA, RunUMAP, FindNeighbors, and FindClusters (*34, 37*). Percentages of cells belonging to each Louvain cluster are shown in Table S15. Normalized data from all samples combined were used for imputation using ALRA with default settings for the purposes of data visualization as shown in Figures 4A-D, S4B-D, and 5C (*75*). Marker gene expression was compared between ALRA-imputed and unimputed data to establish that imputation did not substantially marker gene expression patterns in our dataset (Figure S4, Table S15). Data normalization and integration, UMAP embedding, and Louvain clustering were performed prior to imputation. The threshold for identifying *Gbx2*-positive cells was set as an imputed *Gbx2* expression value greater than 0.1. This threshold was also used for identifying percentages of marker gene-positive cells in unimputed and imputed data as shown in Table S15. All gene expression scaling and centering for visualization purposes was performed on normalized imputed or unimputed data using the Seurat ScaleData function with default parameters (scale.max=10) (*34*).

##### MELD, MAGIC, kNN-DREMI Analyses

Cells belonging to mesenchymal cell clusters (clusters 1-4, see Fig. 4A, C) from all genotypes were used for MELD, MAGIC, kNN-DREMI, and Gene Set Enrichment Analysis (GSEA). Scaled data matrices from the Seurat object integrated assay were loaded using scprep for MELD, MAGIC, and kNN-DREMI (https://github.com/krishnaswamylab/scprep). MELD and MAGIC both denoise scRNA-seq data using graphs to model cellular state space. The same graph signal was used for both MELD and MAGIC as calculated by graphtools.Graph with n_pca=20, decay=40, and knn=10. MELD was run on one-hot vectors for each genotype independently using default parameters (*51*). MAGIC was performed using the same graph signal as MELD (*50*). We used the kNN-DREMI implementation provided in scprep and kNN-DREMI was run on MAGIC-imputed data (*49*). kNN-DREMI analysis was used in order to identify genes with expression levels associated with either *Gbx2* expression in humanized hindlimb or cells with increased humanized RL as calculated using MELD. MAGIC was employed only for the purpose of generating denoised gene expression values for kNN-DREMI analysis of gene-gene relationships but was not used for data visualization, clustering, or sample-associated density estimation using MELD.

#### Skeletal Staining

E18.5 skeletons from two litters from each of *HACNS1* homozygous, chimpanzee ortholog line, and wild type homozygous crosses (n=48 embryos) were stained with Alcian Blue and Alizarin Red as previously described (*65*). Skeletons were imaged under a stereo microscope (Leica S6D). Bone and cartilage lengths of the forelimb and hindlimb pelvic girdle, stylopod, zeugopod, and autopod were measured blinded to genotype using ImageJ 2.0.0. Forelimb measurements include metacarpals 1-5 (cartilage), proximal phalanges 1-5 (cartilage), intermediate phalanges 2-5 (cartilage), distal phalanges 1-5 (cartilage), scapula (bone and cartilage), humerus (bone and cartilage), radius (bone and cartilage), and ulna (bone and cartilage). Hindlimb measurements include metatarsals 1-5 (cartilage), proximal phalanges 1-5 (cartilage), intermediate phalanges 2-5 (cartilage), distal phalanges 1-5 (cartilage), tibia (bone and cartilage), femur (bone and cartilage), pelvis (cartilage), ilium (bone), ischium (bone), pubis (bone), fibula (bone), calcaneum (cartilage), and talus (cartilage). Digit length was calculated as the sum of all metacarpal/metatarsal and phalanx segments. Raw measurements and digit length were normalized to the length of ossified humerus or femur for forelimb or hindlimb digits, respectively. Phalange to metacarpal ratio was calculated as the ratio of the sum of the phalange lengths of each digit to the corresponding metacarpal/metatarsal segment. Interdigital ratios were calculated using raw digit lengths. Raw measurements and images are available at noonan.ycga.yale.edu/noonan_public/Dutrow_HACNS1/.

#### ANOVA Analysis for Gene Expression and Morphometry

ANOVA analysis was performed with the lme4 package in R (RRID: SCR_015654) using default parameters to dissect the effects of genotype on *Gbx2* expression (qRT-PCR data) and limb segment length (morphometric data) (*78*). For the qRT-PCR dataset, we employed an additive model of ΔCt to calculate the main effects of genotype and timepoint and a genotype:timepoint interaction term *(Ct_Gbx2_ – Ct_Hprt1_ ∼ Genotype * Timepoint).* For the morphometric datasets, we calculated the effects of genotype, litter, sex, forelimb versus hindlimb, digit number, and right versus left (RL) on normalized digit length, phalange to metacarpal ratio and interdigital ratio *(Length Ratio ∼ Genotype * (1|Genotype/Litter) * Sex * Limb * Digit * (1|RL) * (1|Litter/Embryo) * (1|Sex/Embryo) * (1|Genotype/Embryo))*. For both datasets, multiple comparisons adjustment was performed using the Benjamini & Hochberg method (*79*).

### Supplemental Note

To obtain further insight into primate-rodent sequence differences at the *HACNS1* locus, we conducted independent global pairwise alignments of the mouse, human and chimpanzee alleles using the EMBOSS Needle tool (Table S1) (*80*). We first considered the entire ∼1.2 kb human or chimpanzee sequence introduced to replace the orthologous endogenous locus in each line. The sequence identity between the human (hg19, chr2:236773456-236774696) and chimpanzee alleles (panTro4, chr2B:241105291-241106530) in the edited mouse lines is 98.2% (22 sequence differences total, of which 15 are fixed in humans). The sequence identity between the human allele and the mouse ortholog replaced in the edited mouse lines (mm9, chr1:91610327-91611486) is 68.6% with 14.5% gapped positions. The sequence identity between the chimpanzee allele and the mouse ortholog is 70.8% with 12.9% gapped positions. The chimpanzee and mouse sequences are identical at 18 of the 22 substituted positions in the human and chimpanzee alleles mentioned above.

*HACNS1* itself, as defined in Prabhakar *et al.* 2006, is highly conserved among sarcopterygian vertebrates (i.e., lobe-finned fishes and tetrapods), including chimpanzee and mouse (*3*). In pairwise alignments, *HACNS1* and its mouse ortholog are 88.6% identical, with 2.2% gapped positions (including potential single nucleotide variants and segregating indels, which we did not exclude in this analysis). The chimpanzee and mouse orthologs are 91.2% identical, with 2.2% gapped positions. Therefore, the majority of the sequence differences between human and mouse, and between chimpanzee and mouse, that we discuss above, fall outside of this core region. It is possible that primate- or rodent-specific sequence differences within the *HACNS1* core interval or in the flanking regions included in our targeting scheme may have contributed to potential primate-specific or rodent-specific changes in the ancestral enhancer function of the *HACNS1* locus.

**Fig. S1.**
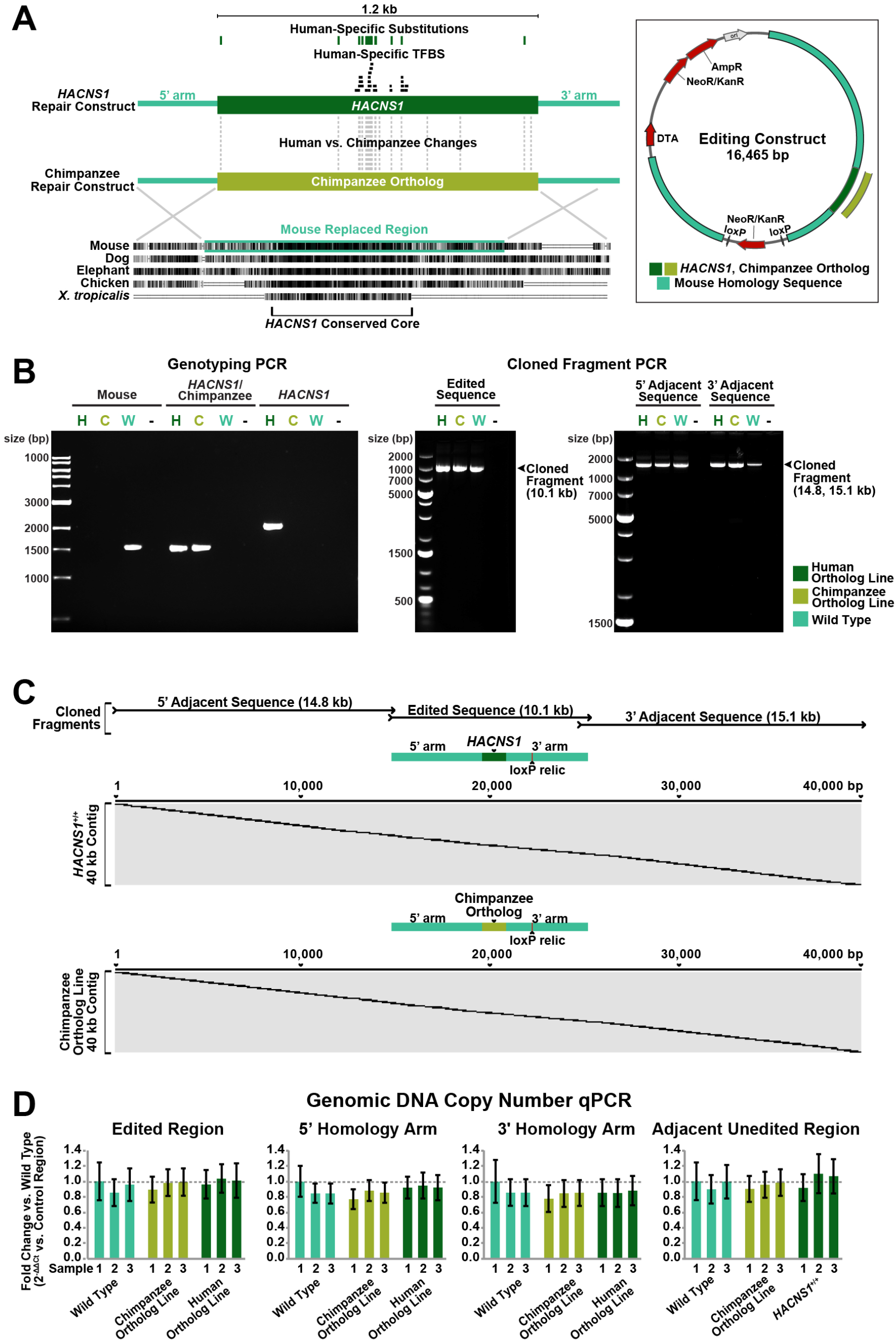
Development and validation of the *HACNS1* and chimpanzee ortholog mouse models. **(A)** *Left: HACNS1* and chimpanzee ortholog line editing constructs are shown with the orthologous replaced region in mouse genome aligned to other vertebrate species derived from the 100-way Multiz alignment in the UCSC hg19 assembly. Non-polymorphic, fixed human-specific substitutions are shown above the human construct and all human versus chimpanzee sequence differences are shown below. The transcription factor binding sites (TFBS) unique to *HACNS1* versus the chimpanzee ortholog shown are based on predictions of JASPAR core mammal motifs in the human and chimpanzee genomes (see also Table S2) (*15*). *Right:* Embryonic stem cell editing construct showing antibiotic resistance (NeoR/KanR), diphtheria toxin (DTA), mouse homology, and location of human or chimpanzee sequences. **(B)** *Left:* PCR products generated with primers specific to the mouse, both *HACNS1* and chimpanzee, and *HACNS1* only orthologs were used for genotyping of *HACNS1* homozygous (labeled as H), chimpanzee ortholog line (labeled as C), and wild type (labeled as W) mice from the F9 or later generation. *Middle:* PCR products were generated using primers outside the homology arms for Sanger sequencing of the edited locus. *Right:* PCR products were generated using primers anchored in the 5’ homology arm and 14.8 kb upstream (5’ adjacent sequence) and primers anchored in 3’ homology arm and 15.1 kb downstream (3’ adjacent sequence) for Sanger sequencing of the regions surrounding the editing locus. **(C)** Sanger sequencing contigs of cloned PCR products from (B), spanning the 40 kb region surrounding edited locus for *HACNS1* homozygous (*top*) and the chimpanzee ortholog line (*bottom*). **(D)** *HACNS1* homozygous, chimpanzee ortholog line, and wild type genomic DNA qPCR for primers specific to the edited region, 5’ homology arm, 3’ homology arm, and adjacent unedited region. All Ct values are normalized to a region on chromosome 5. Three biological replicate samples are shown per genotype. Error bars denote standard deviation between technical replicates.

**Fig. S2.**
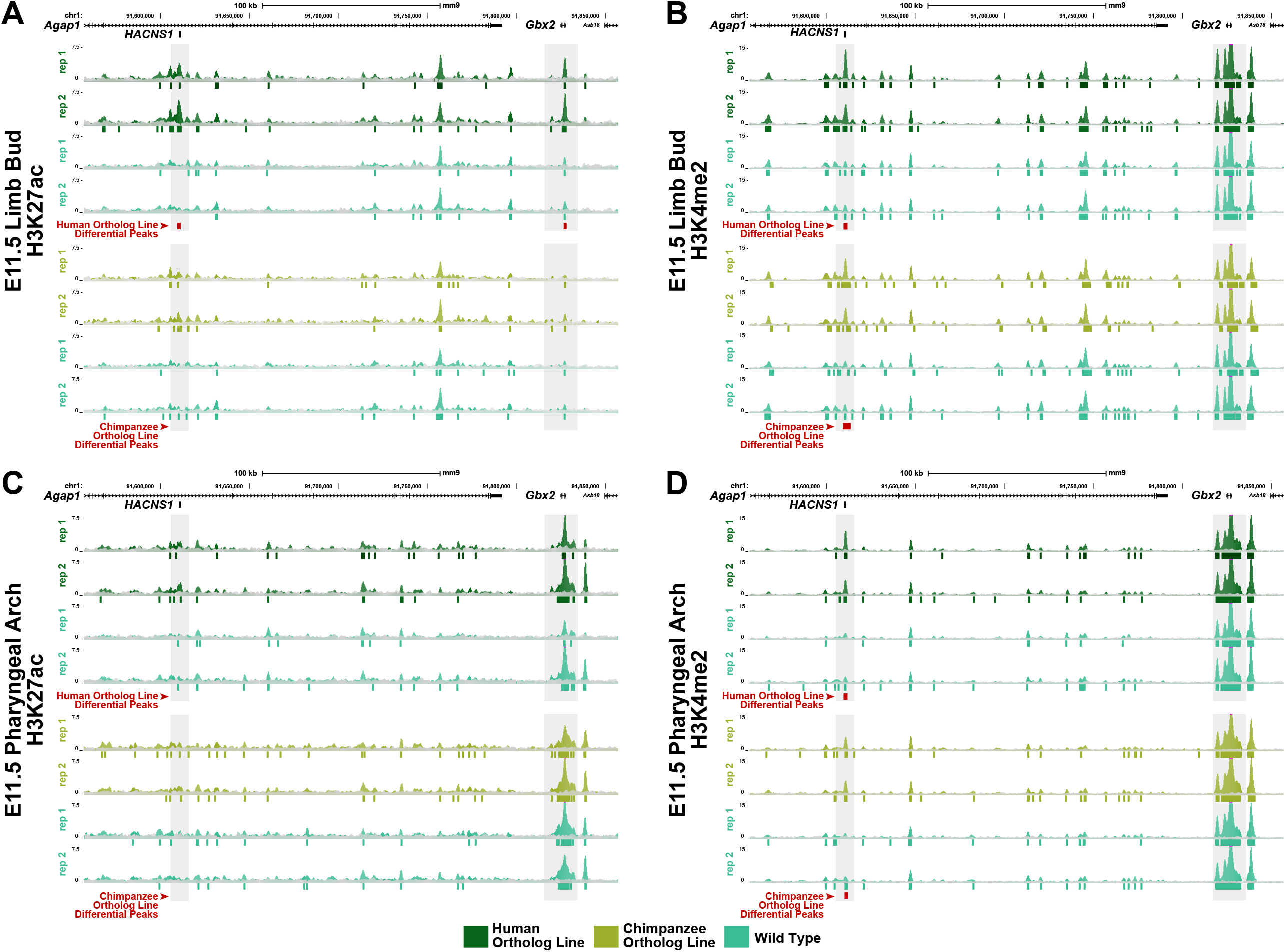
H3K37ac and H3K4me2 ChIP-seq analyses in limb bud and pharyngeal arch. Normalized H3K27ac (*left*) and H3K4me2 (*right*) epigenetic signals in the region spanning the full *HACNS1*-*Gbx2* locus for two biological replicates per genotype for E11.5 limb bud **(A, B)** and pharyngeal arch **(C, D)**. All corresponding input signal tracks are shown overlayed in gray. The location of the edited *HACNS1* locus relative to nearby genes is shown above each track group with a black bar directly below the corresponding UCSC mm9 genome track. *HACNS1* and *Gbx2* loci are highlighted in gray. All significant peaks are represented by genotype-specific colored bars below the signal tracks for *HACNS1* homozygous (in dark green), chimpanzee ortholog line (in olive), and litter-matched wild type (in teal). Peak calls showing significant signal increases between *HACNS1* homozygous and litter-matched wild type, or chimpanzee ortholog line and litter-matched wild type, are shown as red squares below each track group. Detailed differential peak information is available in Table S3.

**Fig. S3.**
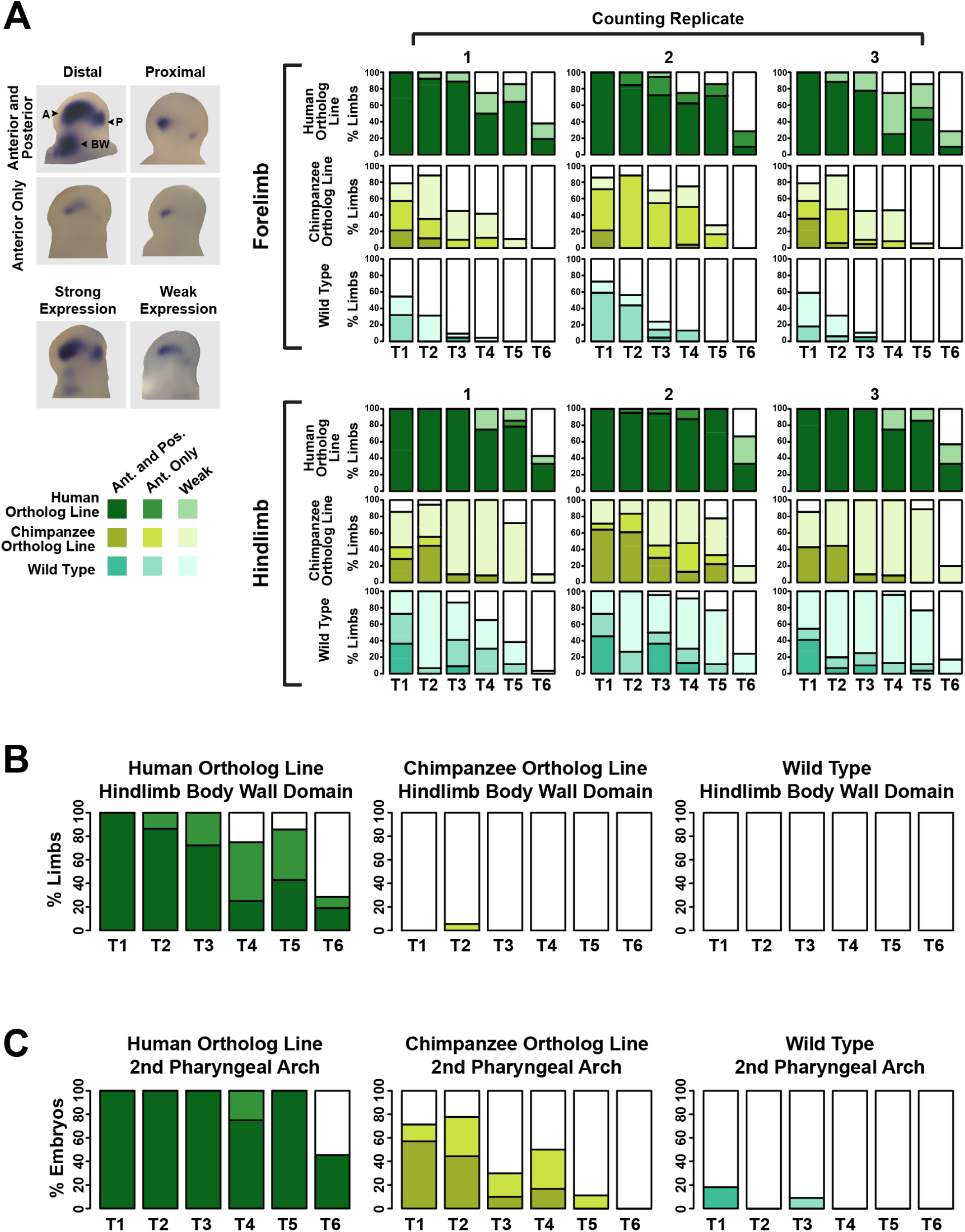
Qualitative analysis of *Gbx2* expression patterns. **(A)** *Left:* representative images of anterior, posterior, proximal, distal (top), and strong versus weak staining patterns (bottom). Anterior (A), posterior (P), and body wall (BW) domains are denoted on top left limb bud. *Right: Gbx2* ISH staining pattern data across 6 developmental timepoints from each of three independent, blinded scorers (marked at top as counting replicates 1-3; see text and Fig. 3A for timepoint scheme). The darkest shade for *HACNS1* homozygous (dark green), chimpanzee ortholog line (olive), and wild type (teal) represents percentage of forelimbs or hindlimbs showing strong anterior and posterior limb bud staining. Medium-dark shade, as shown in legend on the left, denotes strong anterior staining only, while the lightest shade denotes weak staining in any domain. Total numbers of forelimbs and hindlimbs analyzed are shown in Fig. 3B. Photo Credit: Angeliki Louvi, Yale University. **(B)** Counting data for presence or absence of hindlimb bud body wall domain are shown for each genotype. Strong versus weak staining is denoted by darker versus lighter shade, as in (A). **(C)** Counting data for presence or absence of 2^nd^ pharyngeal arch staining domain are shown for each genotype. Strong versus weak staining is denoted by darker versus lighter shade, as in (A).

**Fig. S4.**
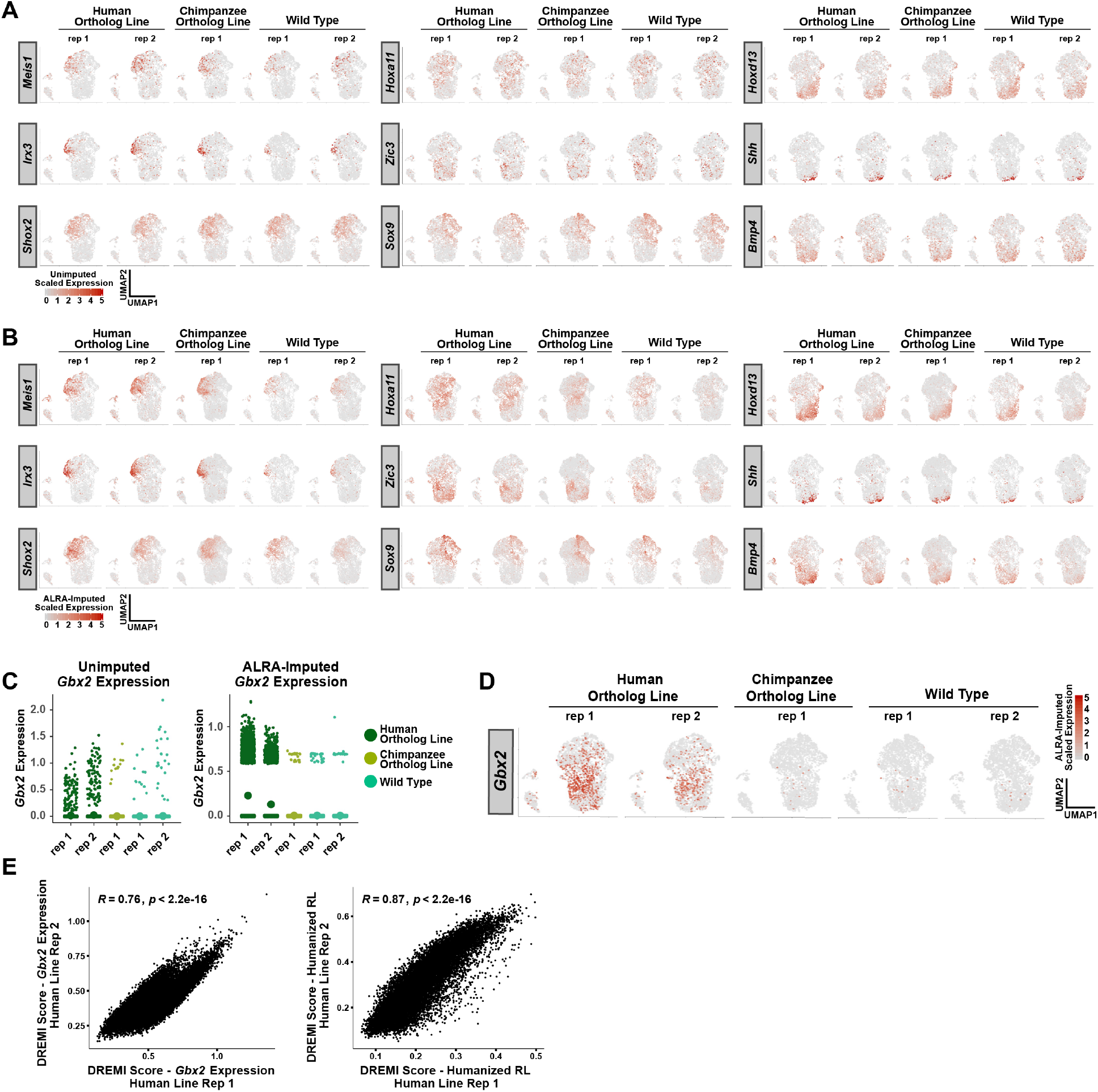
*Gbx2* and developmental marker expression in *HACNS1* homozygous, chimpanzee ortholog line, and wild type E11.5 hindlimb bud replicates. **(A)** UMAP embedding of hindlimb bud cells from *HACNS1* homozygous, chimpanzee ortholog line, and wild type replicates showing conserved expression of representative proximal-distal (*top row*), anterior-posterior (*middle row*), and chrondrogenesis-apoptosis marker genes (*bottom row*). Gene expression data are unimputed and library size-normalized and were centered and scaled using z-scores for plotting (see Materials and Methods) (*75*). See text and Fig. 4B for details on marker genes. **(B)** UMAP embedding of marker gene expression imputed using ALRA and centered and scaled using z-scores) (*75*). See also Table S15. (C) *Left*: Unimputed, library size-normalized *Gbx2* expression values by replicate. *Right*: ALRA-imputed *Gbx2* expression values by replicate. Dots indicate mean expression for each sample. **(D)** UMAP embedding of hindlimb bud cells from *HACNS1* homozygous, chimpanzee ortholog line, and wild type replicates showing *Gbx2* expression values imputed using ALRA and centered and scaled using z-scores (see Materials and Methods) (*75*). **(E)** DREMI scores for association with *Gbx2* (left) or humanized relative likelihood (RL, right) for genes expressed in humanized replicate 2 versus replicate 1. Spearman correlation values are shown for each plot with associated p values.

**Fig. S5.**
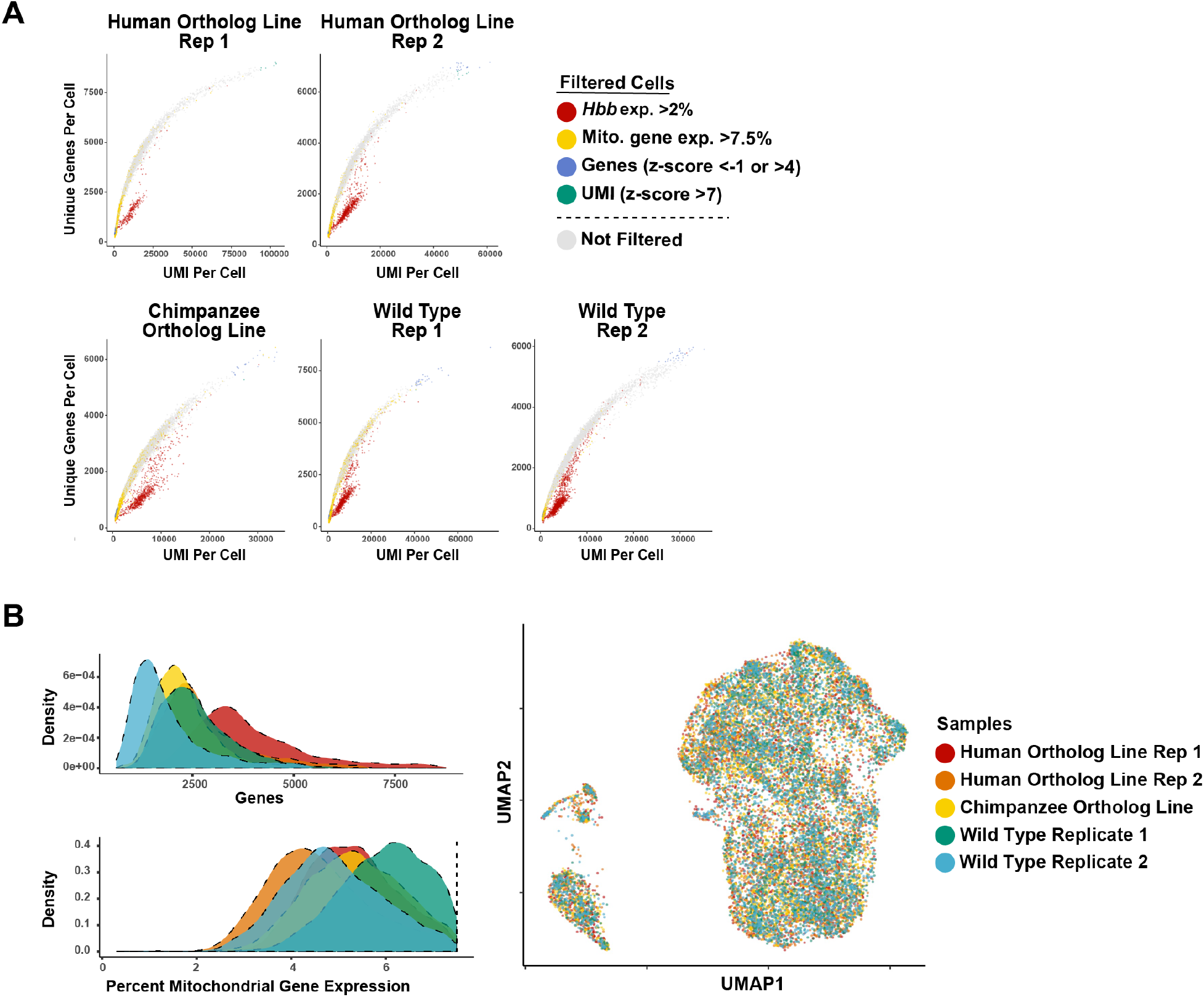
Sample quality metrics in *HACNS1* homozygous, chimpanzee ortholog line, and wild type E11.5 hindlimb bud replicates. **(A)** UMI per cell versus number of unique genes per cell for *HACNS1* homozygous, chimpanzee ortholog line, and wild type scRNA-seq replicates before filtering. Points representing cells are colored by the indicated filtering criteria. **(B)** *Left*: Density plots showing genes per cell (top) and percent mitochondrial gene expression per cell (bottom) for each replicate after filtering. Dashed line indicates 7.5% mitochondrial DNA expression cutoff implemented in filtering. *Right:* UMAP embedding showing each replicate colored separately.

**Fig. S6.**
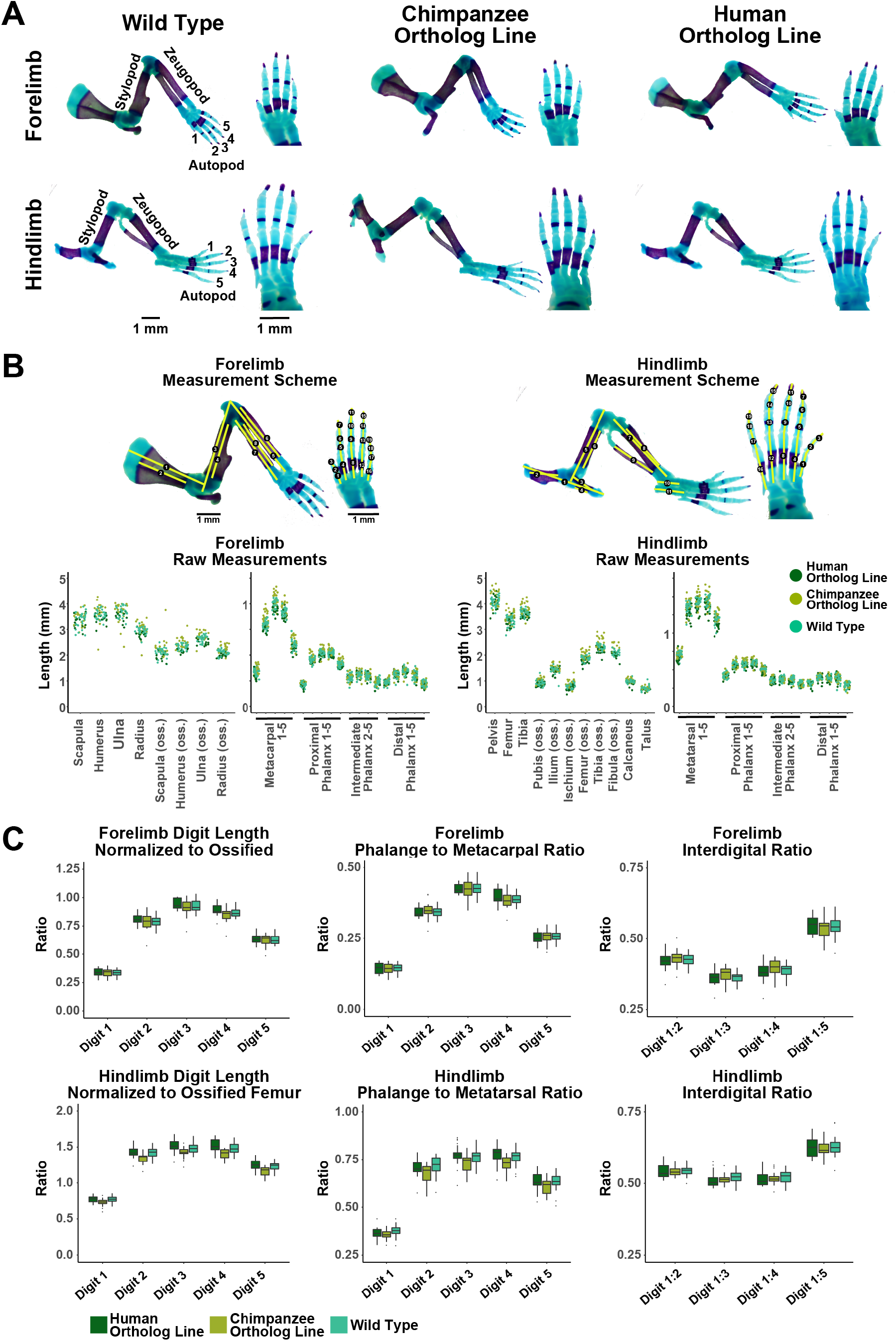
Morphometric analyses of *HACNS1* homozygous, chimpanzee ortholog line, and wild type skeletons. **(A)** Representative images of E18.5 forelimbs and hindlimbs of the indicated genotypes stained with Alizarin Red (violet; bone) and Alcian Blue (blue; cartilage). Forelimb and hindlimb autopod, zeugopod, and stylopod are shown on the left, and numbers indicate digit identity. High magnification images of forelimb and hindlimb autopods are shown on the right. Photo Credit: Emily Dutrow, Yale University. **(B)** *Top:* Measurement scheme for E18.5 skeleton morphometric analysis is shown with yellow lines denoting measured segments for forelimb zeugopod and stylopod cartilage and/or bone: scapula (1,2), humerus (3,4), ulna (5,6), radius (7,8); hindlimb zeugopod and stylopod cartilage and/or bone: pelvis (1), ilium (2), pubis (3), ischium (4), femur (5,6), tibia (7,8), fibula (9), talus (10), calcaneus (11), and metacarpal/metatarsal and phalange autopod segments (digit 1: 1-3; digit 2: 4-7; digit 3: 8-11; digit 4: 12-15; digit 5:16-19). *Bottom:* Raw data for all measured segments are plotted and colored by genotype. **(C)** Normalized digit length, phalange to metacarpal/metatarsal ratio, and interdigital ratio for forelimb and hindlimb digits are shown by genotype. Digit length is calculated as sum of all metacarpal/metatarsal and phalange segments. Forelimb and hindlimb digit lengths are normalized to ossified humerus and femur length of the same sample digit length, respectively. For ANOVA analysis of morphometric data see Tables S9-S11.

